# *Mycobacterium tuberculosis* PhoP integrates stress response to intracellular survival by regulating cAMP level

**DOI:** 10.1101/2022.03.07.483216

**Authors:** Hina Khan, Partha Paul, Harsh Goar, Bhanwar Bamniya, Navin Baid, Dibyendu Sarkar

## Abstract

Survival of *M. tuberculosis* within the host macrophages requires the bacterial virulence regulator PhoP, but the underlying reason remains unknown. cAMP is one of the most widely used second messengers, which impacts on a wide range of cellular responses in microbial pathogens including *M. tuberculosis*. Herein, we hypothesized that intra-bacterial cAMP level could be controlled by PhoP since this major regulator plays a key role in bacterial responses against numerous stress conditions. A transcriptomic analysis reveals that PhoP functions as a repressor of cAMP-specific phosphodiesterase (PDE) Rv0805, which hydrolyses cAMP. In keeping with these results, we find specific recruitment of the regulator within the promoter region of *rv0805* PDE, and absence of *phoP* or ectopic expression of *rv0805* independently accounts for elevated PDE synthesis leading to depletion of intra-bacterial cAMP level. Thus, genetic manipulation to inactivate PhoP-*rv0805*-cAMP pathway decreases cAMP level, stress tolerance and intracellular survival of the bacillus.

## Introduction

*M. tuberculosis*, the causative agent of pulmonary tuberculosis, encounters diverse environmental conditions during infection, persistence and transmission of the disease (Ehrt *et al*, 2018; Ernst, 2012; Russell, 2011). However, the pathogen is exceptionally capable to adjust and survive within diverse host environments. Mycobacterial adaptive response to different stages of infection is achieved via fine tuning of regulation of gene expression using an extensive repertoire of more than 100 transcriptional regulators, 11 two-component systems, 6-serine-threonine protein kinases and 13 alternative sigma factors suggesting that a very complex transcriptional program is important for *M. tuberculosis* pathogenesis. While regulation of gene expression as a consequence of interaction of tubercle bacilli with its immediate environment remains critically important (Rohde *et al*, 2007), defining these regulatory pathways represent a major challenge in the field.

3’, 5’-cyclic adenosine monophosphate (cAMP), one of the most widely used second messengers, impacts on a wide range of cellular responses in microbial pathogens including *M. tuberculosis* (McDonough & Rodriguez, 2011). The bacterial genome encodes at least 15 adenylate cyclases, including one of the adenylate cyclases (Rv0386) which is required for virulence (Agarwal *et al*, 2009), and multiple cAMP regulated effector proteins(Banerjee *et al*, 2015; Johnson & McDonough, 2018; McDonough & Rodriguez, 2011; Shenoy & Visweswariah, 2006). cAMP levels are elevated upon infection of macrophages by pathogenic mycobacterium (Bai *et al*, 2009) and addition of exogenous cAMP was shown to influence mycobacterial protein expression (Gazdik & McDonough, 2005). While intra-bacterial cAMP responds to macrophage environment and carries out specific functions like gene expression and protein function under host-related conditions (Bai *et al*, 2011; Bai *et al*., 2009; Johnson *et al*, 2017; Johnson & McDonough, 2018; Knapp & McDonough, 2014), secreted cAMP controls macrophage signalling (Agarwal *et al*., 2009; Agarwal *et al*, 2006; Gazdik *et al*, 2009; Johnson *et al*., 2017; Nambi *et al*, 2013; Ranganathan *et al*, 2016) (Rittershaus *et al*, 2018). cAMP signalling remains essential to *M. tuberculosis* pathogenesis. Agarwal and colleagues had shown that a burst in synthesis of cAMP upon infection of macrophages, improved bacterial survival by interfering with host signalling pathways (Agarwal *et al*., 2009). In keeping with this, anti-tubercular compounds that interfere with mycobacterial cAMP levels also impact intracellular growth of mycobacteria (Bai *et al*., 2011; Johnson *et al*., 2017; Knapp *et al*, 2015; Nambi *et al*., 2013; Rittershaus *et al*., 2018; VanderVen *et al*, 2015; Wilburn *et al*, 2022). Together, these results underscore the significance of cAMP signalling in mycobacteria.

cAMP signalling is controlled at the transcriptional level. Previous *in silico* studies had identified two out of 10 predicted nucleotide binding proteins of *M. tuberculosis* as members of the CRP/FNR superfamily of transcriptional regulators (McCue *et al*, 2000). These are CRP (cyclic AMP receptor protein), encoded by *rv3676* and CMR (cAMP macrophage regulator), encoded by *rv1675c*, respectively. Of these two, CRP becomes activated upon cAMP binding, and functions as a global regulator of ∼100 genes (Agarwal *et al*., 2006; Bai *et al*, 2005; Rickman *et al*, 2005). Consequently, deletion of the *crp* locus significantly impairs mycobacterial growth and attenuates virulence of the bacilli in a mouse model (Rickman *et al*., 2005). In contrast, CMR is necessary for regulated expression of genes involved in virulence and persistence including members of the dormancy regulon (Gazdik *et al*., 2009; Ranganathan *et al*., 2016; Smith *et al*, 2017). These results strongly suggest that cAMP homeostasis i.e. a balance between cAMP synthesis by adenylate cyclases and cAMP degradation by phosphodiesterases contributes to rapid adaptive response of mycobacteria in a hostile intracellular environment (Johnson & McDonough, 2018; McDonough & Rodriguez, 2011). However, very little is known about the underlying mechanisms of regulation of mycobacterial cAMP level.

*M. tuberculosis* encounters a hostile environment within the host. Among the hostile conditions is the acidic pH stress and exposure to host immune effectors such as NO (Nathan & Shiloh, 2000; Rustad *et al*, 2009; Wayne & Sohaskey, 2001). A growing body of evidence connects virulence-associated mycobacterial *phoP* locus with varying environmental conditions, including acid stress (Abramovitch *et al*, 2011; Bansal *et al*, 2017; Tan *et al*, 2013), heat-shock (Sevalkar *et al*, 2019; Singh *et al*, 2014) and integration of acid stress response to redox homeostasis (Baker *et al*, 2019; Baker *et al*, 2014; Goar *et al*, 2022). Disruption of *phoP,* the gene encoding the response regulator of the PhoPR two-component signal transduction system (Gupta *et al*, 2006) significantly reduces *in vivo* multiplication of the bacilli (Walters *et al*, 2006), suggesting that PhoPR remains essential for virulence (Perez *et al*, 2001). Moreover, a mutant lacking this system shows a significantly lowered synthesis of cell-wall components diacyltrehaloses, polyacyltrehaloses, and sulfolipids, specific to pathogenic mycobacterial species (Gonzalo Asensio *et al*, 2006a; Goyal *et al*, 2011; Walters *et al*., 2006). In fact, the significant attenuation of *phoPR* deletion strain forms the basis of the mutant being considered in trials as a vaccine strain (Arbues *et al*, 2013). Recent transcriptomic analyses revealed that approximately 2% of the H37Rv genome is regulated by PhoP (Solans *et al*, 2014; Walters *et al*., 2006), and mycobacterial gene expression in response to acidic pH significantly overlaps with the PhoP regulon (Rohde *et al*., 2007). Consistent with this, a large subset of PhoPR-regulated low-pH-inducible genes are induced immediately following *M. tuberculosis* phagocytosis and remain induced during macrophage infection (Gonzalo Asensio *et al*, 2006b; Martin *et al*, 2006; Perez *et al*., 2001; Walters *et al*., 2006). Additional evidences coupled with more recent results suggest that during onset of macrophage infection, PhoPR activation is linked to acidic pH and the available carbon source, suggesting a physiological link between pH, carbon source, and macrophage pathogenesis (Baker *et al*., 2019).

In this study, we hypothesized that intra-mycobacterial cAMP level could be determined by PhoP since the major regulator has been implicated in regulation of bacterial responses against numerous stress conditions, many of which function as signals to activate cAMP synthesizing diverse adenylate cyclases (Knapp & McDonough, 2014). Our results connect virulence regulator PhoP with intra-mycobacterial cAMP level. We discovered that PhoP regulates expression of cAMP-specific phosphodiesterase *rv0805*, which hydrolyses cAMP. To further probe the regulation, we demonstrate that under the condition which activates PhoP-PhoR system, PhoP in a PhoR-dependent manner represses transcription of *rv0805* through direct DNA binding at the upstream regulatory region.

These observations account for a consistently lower level of cAMP in a PhoPR-deleted *M. tuberculosis* H37Rv (*phoPR*-KO) relative to the WT bacilli, and establishes the molecular mechanism of regulation of cAMP level, absence of which strikingly impacts phagosome maturation, and reduces mycobacterial survival within macrophages and mice. Together, the newly identified mechanism of regulation of cAMP level allows intra-phagosomal survival and growth program of mycobacteria.

## RESULTS

### Intra-mycobacterial cAMP level is regulated by the *phoP* locus

We compared cAMP levels of WT and *phoPR*-KO mutant (lacking both the single copies of *phoP* and *phoR* genes), grown under normal, NO stress and acid stress conditions (Fig. 1A). *phoPR*-KO showed a significantly lower level of cAMP relative to the WT bacilli, both under normal and stress conditions. Complementation of the mutant (Compl.) could restore cAMP to the WT level. A higher cAMP level in the complemented strain under NO stress is possibly attributable to reproducibly higher *phoP* expression in the complemented mutant under specific stress conditions (Khan et al., 2022). Because bacterial growth often varies under stress conditions, and growth inhibition can influence cAMP level, we compared viability of mycobacterial strains under normal and indicated stress conditions (conditions of cAMP measurements) by determining the bacterial CFU (Figure 1 - figure supplement 1). Note that for *in vitro* viability under specific stress conditions, indicated mycobacterial strains were grown to mid log phase (OD_600_ 0.4-0.6) and exposed to acidic media (7H9 media, pH 4.5(Gouzy *et al*, 2021) for further two hours at 37°C. Likewise, for NO stress, cells grown to mid-log phase were exposed to 0.5 mM DetaNonoate for 40 minutes. Our results suggest that WT and *phoPR*-KO under carefully controlled stress conditions display comparable viability, indicating that variation in viable cell counts of the mutant under specific stress conditions do not account for lower cAMP level. From these results, we conclude that PhoP plays a major role in maintaining intra-mycobacterial cAMP level.

**Figure 1:**
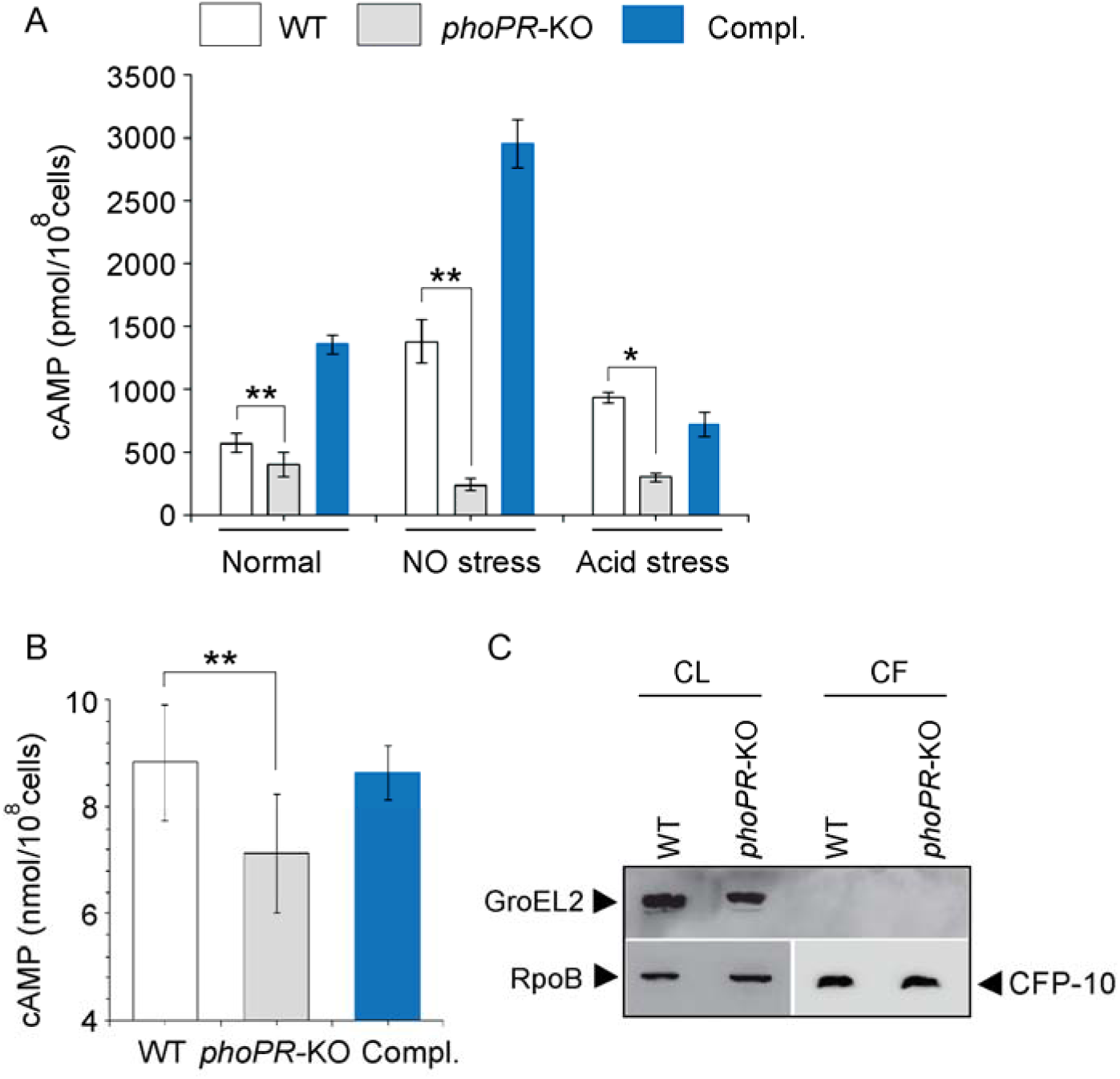
PhoP contributes to maintenance of mycobacterial cAMP level. (A) Intra-mycobacterial cAMP levels were determined by a fluorescence-based assay as described in the Methods, and compared for indicated mycobacterial strains, grown under normal or specific stress conditions. For acid stress, mycobacterial strains were initially grown to mid-log phase (OD_600_ 0.4 to 0.6), and then it was transferred to acidic pH (7H9 media, pH 4.5) for further 2 hours of growth at 37°C. For NO stress, cells grown to mid-log phase were exposed to 0.5 mM DataNonoate for 40 minutes. The data represent average values from three biological repeats (*P≤0.05, **P≤0.01). (B) To compare secretion of cAMP by WT and *phoPR*-KO, cAMP levels were also determined in the corresponding culture filtrates (CF). (C) Immunoblotting analysis of 10 µg of cell lysates (CL) and 20 µg of culture filtrates (CF) of indicated *M. tuberculosis* strains. α-GroEL2 was used as a control to verify cytolysis of cells, CFP-10 detected as a secreted mycobacterial protein in the culture filtrates, and RpoB used as the loading control.

To investigate the possibility that *phoPR*-KO secretes out more cAMP and therefore, shows a lower cytoplasmic level, we compared cAMP secretion of WT and the mutant (Fig. 1B). The WT and mutant strains were grown as described in the Methods and culture filtrates (CF) as well as cell lysates (CL) were collected as described previously (Anil Kumar *et al*, 2016). Our results demonstrate that the mutant reproducibly secretes lower amount of cAMP relative to the WT bacilli, and cAMP secretion is fully restored in the complemented mutant (Compl.). As the fold difference of secretion (∼ 1.25-fold) is not much different relative to the fold difference in intra-mycobacterial cAMP level (∼2-fold), we suggest that lower cAMP level of the mutant is not due to its higher efficacy of cAMP secretion. Fig. 1C confirms absence of autolysis of mycobacterial cells as GroEL2, a cytoplasmic protein, was undetectable in the culture filtrates (CF).

### PhoP functions as a repressor of *rv0805*

We next investigated role of the *phoP* locus on expression of mycobacterial adenylate cyclases (ACs) and phosphodiesterases (PDEs), which synthesize and degrade cAMP, respectively (Fig. 2A). The selection of ACs and PDEs were based on two key points. First, we have chosen ACs which are activated by known signals (Knapp & McDonough, 2014). Second, we reasoned that the previously reported ACs were activated under environmental conditions, which are linked to mycobacterial *phoP* locus (Bansal *et al*., 2017; Goar *et al*., 2022). Our RT-qPCR results using gene-specific primer pairs (Supplementary file 1a) suggest that expression of adenylate cyclases including *rv0386*, *rv1264*, *rv1647* and *rv2488c* (Agarwal *et al*., 2009; Dass *et al*, 2008; Dittrich *et al*, 2006; Knapp & McDonough, 2014) do not appear to be regulated by the *phoP* locus. However, a significant activation of adenylate cyclase *rv0891c* (4±0.05-fold), and repression of phosphodiesterase *rv0805* expression (6.5±0.7-fold), respectively, were dependent on the *phoP* locus. Although Rv0891c was suggested as one of the *M. tuberculosis* H37Rv adenylate cyclases, the protein lacks most of the important residues conserved for adenylate cyclase family of proteins (Zaveri *et al*, 2021). On the other hand, *rv0805* encodes for a cAMP specific PDE, present only in slow growing pathogenic *M. tuberculosis* (Matange *et al*, 2013; Shenoy *et al*, 2007; Shenoy *et al*, 2005). Although expression of *rv0805* was restored at the level of WT in the complemented mutant (Compl.), we observed poor restoration of expression of *rv0891c* in the Compl. strain. Further, expression of PDE *rv1339* which contributes to mycobacterial cAMP level (Thomson *et al*, 2022) remains unaffected by the *phoP* locus. Therefore, we focussed our attention on the biological significance of PhoP-dependent regulation of *rv0805*. It should be noted that expressions of *rv1357c* and *rv2837c*, encoding PDEs for cyclic di-GMP (Flores-Valdez *et al*, 2015) and cyclic di-AMP (Valadares & Woo, 2017), respectively, remained unchanged in *phoPR*-KO.

**Figure 2:**
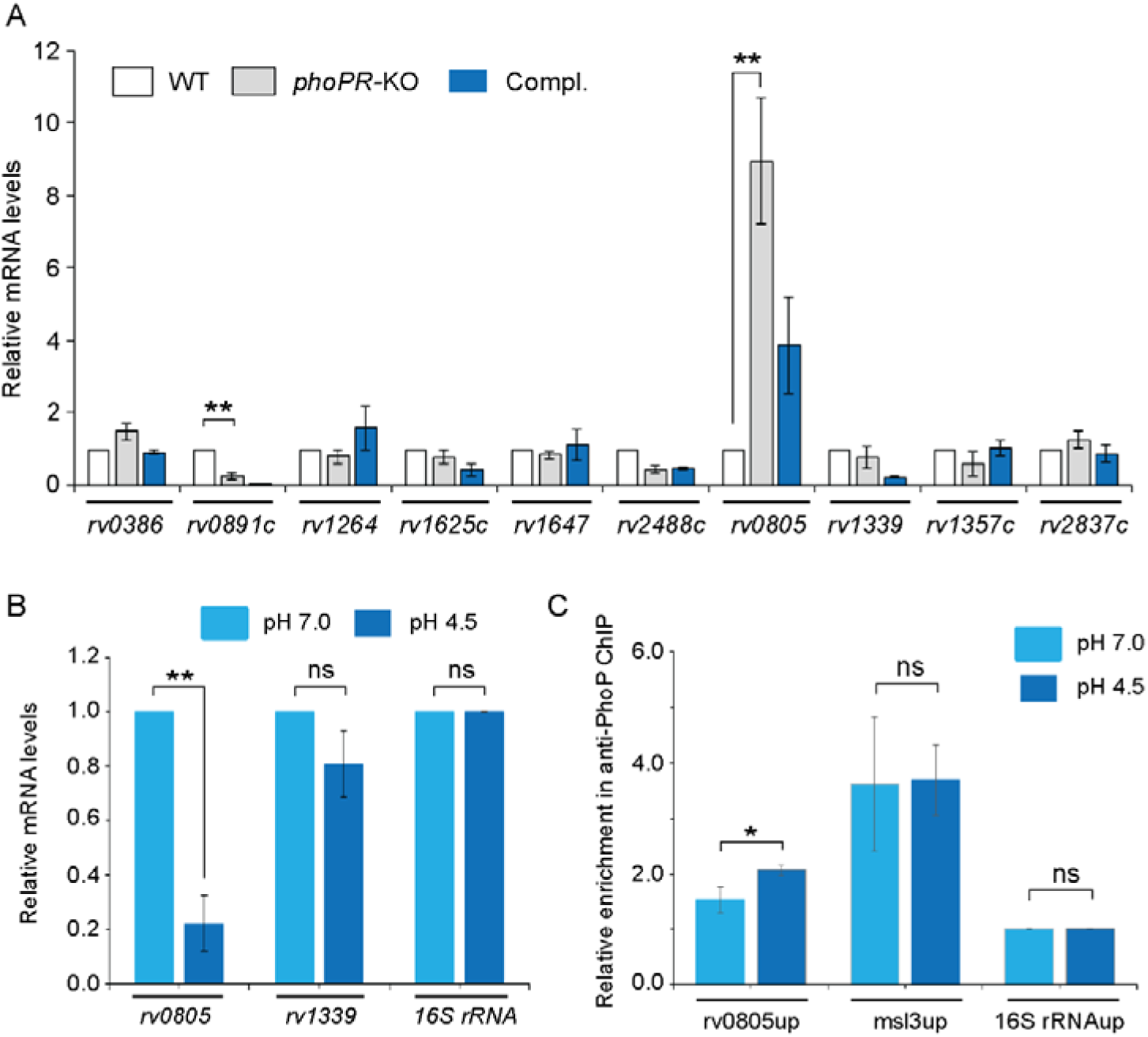
PhoP regulates expression of phosphodiesterase (PDE) *rv0805*. (A) To investigate regulation of cAMP level, mRNA levels of well-characterized adenylate cyclases, and PDEs were compared in indicated mycobacterial strains by RT-qPCR as described in the Methods. The results show average values from biological triplicates, each with two technical repeats. Note that the difference in expression levels of *rv0805* between WT and *phoPR*-KO was significant (p<0.01), whereas the fold difference in mRNA level between WT and the complemented mutant (Compl.) remains nonsignificant (not indicated). (B) To determine the effect of acidic pH conditions of growth, mycobacterial *rv0805* expression was compared in WT grown under normal (pH 7.0) and acidic pH (pH 4.5). Average fold difference in mRNA levels from biological duplicates (each with a technical repeat) were measured as described in the Methods (**P≤0.01). As controls, expression of *rv1339* and *16S rDNA* were also measured. Non-significant difference is not indicated. (C) *In vivo* PhoP binding to *rv0805* promoter (rv0805up) was compared in WT grown under normal and acidic conditions of growth using anti-PhoP antibody followed by ChIP-qPCR. Fold enrichment data represent mean values of two independent experiments with a statistically significant fold difference (\*\**P*-value <0.01; \**P*-value <0.05). The upstream regulatory regions of 16S rRNA (16S rRNAup) and msl3 (msl3up) were used as negative and positive controls, respectively. The assay conditions, sample analyses, and detection are described in the Methods section.

As we reproducibly observed activation of *rv0805* expression in *phoPR*-KO (relative to WT), we investigated whether acidic pH conditions under which *phoPR* system is activated (Abramovitch *et al*., 2011; Bansal *et al*., 2017), impacts expression of *rv0805* in WT bacilli (Fig. 2B). Our results show that repression of *rv0805* is significantly higher in WT grown under acidic conditions (pH 4.5) relative to normal conditions (pH 7.0) of growth. This observation is consistent with RNA-seq data displaying significant down-regulation of *rv0805* in WT bacilli grown under acidic pH conditions relative to normal conditions of growth (GEO Accession number: GSE180161). As a control, expression of PDE *rv1339* which also degrades cAMP, remains unaffected under acidic conditions of growth. The finding that acidic pH (pH4.5) conditions of growth promoted PhoP-dependent repression of *rv0805*, prompted us to investigate whether PhoP directly binds to *rv0805* promoter. To this end, we next examined *in vivo* recruitment of PhoP within the *rv0805* promoter by chromatin immunoprecipitation (ChIP) experiments (Fig. 2C). In this assay, formaldehyde-cross-linked DNA-protein complexes of growing *M. tuberculosis* cells were sheared to generate fragments of average size ≍500 bp. Next, ChIP experiments utilized anti-PhoP antibody, and IP DNA was analysed by real-time qPCR relative to a mock sample (without antibody as a control) using FPrv0805up/RPrv0805up as the primer pair (Supplementary file 1a). Our results show that under normal condition (light blue bars) rv0805up showed an insignificant enrichment of PCR signal for PhoP relative to mock (no antibody control) sample, suggesting low affinity DNA binding of PhoP under normal conditions.

However, IP samples from cells grown under acidic pH showed a significantly higher enrichment of PhoP at the *rv0805* promoter (rv0805up; compare *light blue* bars with *dark blue* bars). As controls, promoter of *msl3* (msl3up) which is controlled by PhoP, and non-specific 16S rRNAup showed comparable enrichment, and no enrichment under normal and acidic conditions, respectively. Thus, ChIP data showing PhoP recruitment under acidic pH conditions is in agreement with low pH-specific impact of PhoP on *rv0805* expression (Fig. 2B). Note that PhoP binding to msl3up was used as a positive control.

To examine DNA binding *in vitro*, we first probed for the PhoP binding site within the upstream regulatory region of *rv0805* (rv0805up) by MEME Bioinformatic software using the consensus PhoP binding sequence (He & Wang, 2014). Our results suggest that the likely PhoP binding sequence spans from -127 to -110 (relative to the ORF start site) of rv0805up (Figure 2-figure supplement 1A) (p=0.000726). Next, recombinant PhoP was phosphorylated by acetyl -phosphate (AcP) and used in EMSA experiments as described earlier (Pathak *et al*, 2010). Consistent with the presence of a PhoP binding site, EMSA results demonstrate that P∼PhoP binds to radio-labelled rv0805up to form a complex stable to gel electrophoresis (Figure 2-figure supplement 1B).

### Probing PhoP-dependent regulation of *M. tuberculosis rv0805*

To examine whether the regulatory effect was attributable to PhoP activation via phosphorylation, we next grew *phoPR*-KO complemented with either *phoP* (Fig. 3A) or the entire *phoPR* encoding ORFs (Fig. 3B), both under normal (pH 7.0; empty bars) and acidic (pH 4.5; black bars) conditions and compared relative expression of *rv0805*. Although both strains expressed *phoP*, the former strain lacked a functional copy of *phoR*, the cognate sensor kinase which phosphorylates PhoP (Gupta *et al*., 2006). *M. tuberculosis* H37Rv lacking a *phoR* gene (*phoPR*-KO*::phoP*) did not show a low pH-dependent repression of *rv0805* expression. However, similar to WT bacilli, we observed a low pH-dependent significant down-regulation of *rv0805* expression in *phoPR*-KO*::phoPR* (Compl.). Note that a comparable expression of PDE *rv1339* was observed in both strains regardless of growth conditions. These results indicate that acidic pH-dependent repression of *rv0805* expression *in vivo* is attributable to P∼PhoP requiring the presence of PhoR.

**Figure 3:**
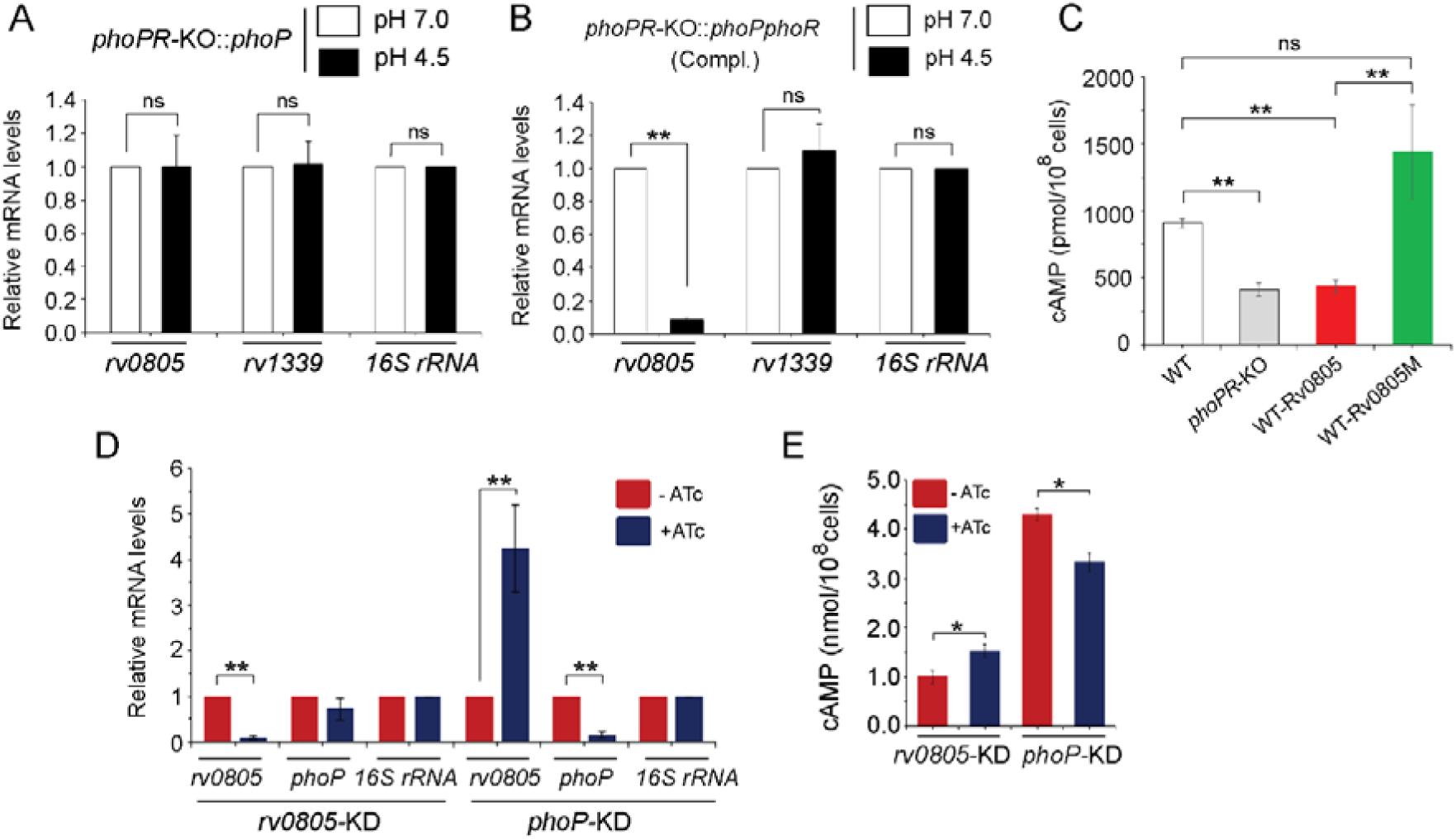
PhoP dependent repression of *rv0805* to maintain mycobacterial cAMP level requires the presence of PhoR. (A-B) To determine the impact of PhoR (the cognate sensor kinase of PhoP), expression of *rv0805* was compared in indicated *M. tuberculosis* H37Rv strains: (A) *phoPR*-KO*::phoP* (*phoPR* mutant complemented with *phoP*) and (B) *phoPR*-KO*::phoPR* (*phoPR* mutant complemented with *phoP-phoR*) under normal and acidic conditions of growth. As expected, *phoPR*-KO*::phoPR* (Compl.) shows a significant repression of *rv0805* (but not *rv1339*) under acidic pH (***P<0.001). However, *rv0805* expression remains comparable in *phoPR*-KO*::phoP* under normal as well as acidic conditions of growth. As a control, *rv1339* fails to show a variable expression in indicated mycobacterial strains. (C) To determine the effect of ectopic expression of *rv0805* on intra-mycobacterial cAMP level, WT and mutant Rv0805 proteins (Rv0805M, defective for phosphodiesterase activity) were expressed in *M. tuberculosis* H37Rv (to construct WT-Rv0805, and WT-Rv0805M, respectively) as described in the Methods section. Similar to *phoPR*-KO, WT-Rv0805 (but not WT-Rv0805M) showed a considerably lower level of cAMP relative to WT bacteria. Significance in variation of cAMP levels were determined by paired student’s t-test (**P<0.01). (D-E) Relative expression of *phoP* and PDE in *phoP*-KD and *rv0805*-KD (*phoP* and *rv0805* knock-down constructs, respectively). In keeping with elevated expression of *rv0805* in *phoPR*-KO, *phoP*-KD shows an elevated expression of *rv0805* relative to WT bacilli. In contrast, *phoP* expression level remains unaffected in *rv0805*-KD mutant. Panel E measured corresponding intra-bacterial cAMP levels in the respective knock-down mutants, as described in the legend to Fig. 1A.

To examine the effect of Rv0805 on mycobacterial cAMP level, we next expressed a copy of *rv0805* in WT bacteria (referred to as WT***-***Rv0805) (Fig. 3C). *rv0805* ORF was cloned within the multicloning site of pSTki (Parikh *et al*, 2013) between the EcoRI and HindIII sites under the control of P_myc1_*tetO* promoter, and expression of *rv0805* under non-inducing condition was verified by determining the mRNA level (Figure 3 - figure supplement 1A). Although copy number of episomal vectors with pAl5000 origin of replication (*oriM*) have been reported to be 3 by Southern hybridization (Ranes *et al*, 1990), in this case wild-type and mutant Rv0805 proteins were expressed from single-copy chromosomal integrants (Parikh *et al*., 2013). We then assessed the impact of Rv0805 on intra-mycobacterial cAMP level (Fig. 3C). Consistent with a previous study (Agarwal *et al*., 2009), WT-Rv0805 showed a significant depletion (2.1± 0.7-fold) of intra-mycobacterial cAMP relative to WT bacteria. To confirm that the reduced level of mycobacterial cAMP is attributable to *rv0805* expression, we also expressed *rv0805M*, a mutant Rv0805 lacking phosphodiesterase activity. As structural data coupled with biochemical evidences reveal that Asn-97 at the enzyme active site plays a key role in phosphodiesterase activity of Rv0805 (Shenoy *et al*., 2007; Shenoy *et al*., 2005), the mutant Rv0805M was constructed by changing the conserved Asn-97 to Ala. WT-Rv0805M showed an insignificant variation of cAMP level relative to WT, suggesting that depletion of intra-mycobacterial cAMP in WT-Rv0805 is indeed attributable to phosphodiesterase activity of Rv0805. The corresponding mRNA levels of PDEs (wild-type and the mutant) are over-expressed approximately 4.5-6 -fold relative to the genomic *rv0805* level of WT-H37Rv (Figure 3-figure supplement 1A). In contrast, other PDE encoding genes (*rv1357* and *rv2387*), under identical conditions, demonstrate comparable expression levels in WT-H37Rv and *rv0805* over-expressing strains. Over-expression of these PDEs did not influence bacterial growth under normal conditions (Figure 3 - figure supplement 1B).

To further probe regulation of Rv0805 expression and its control of intra-mycobacterial cAMP level, we utilized a previously reported CRISPRi based approach (Singh *et al*, 2016) to construct *rv0805* and *phoP* knockdown (*rv0805*-KD, and *phoP-KD*, respectively) mutants. Consistent with *phoPR*-KO, *phoP*-KD shows a significantly higher *rv0805* expression in the presence of ATc relative to its absence (Fig. 3D). However, despite a significant down regulation of *rv0805* expression in presence of ATc, a comparable *phoP* expression was observed in *rv0805*-KD mutant both in the absence or presence of ATc. As a control, we observed a comparable expression of 16S rRNA in both knock-down mutants. Next, we determined intra-mycobacterial cAMP of the mutants as described in Fig. 1 (Fig. 3E). cAMP level of *phoP*-KD (showing activation of Rv0805) was significantly lower relative to WT bacteria. In contrast, *rv0805*-KD mutant demonstrated a significantly higher level of cAMP relative to WT. We speculate that effective knocking down of *phoP* or *rv0805* is not truly reflected in the extent of variation of cAMP levels possibly due to the presence of numerous other mycobacterial PDEs. These data represent an integrated view of our results suggesting that PhoP-dependant repression of *rv0805* regulates intra-mycobacterial cAMP level. In keeping with these results, activated PhoP under acidic pH conditions significantly represses *rv0805*, and intracellular mycobacteria most likely utilizes a higher level of cAMP to effectively mitigate stress for survival under hostile environment including acidic pH of the phagosome.

### PhoP contributes to mycobacterial stress tolerance by repressing the *rv0805* PDE expression

To investigate whether cAMP level influences mycobacterial susceptibility to stress, we compared *in vitro* growth under acidic pH (pH 4.5) (Fig. 4A). As expected, *phoPR*-KO showed a significant growth inhibition relative to WT under low pH (pH 4.5) (Bansal *et al*., 2017). WT*-* Rv0805, but not WT*-*Rv0805M displayed a comparable susceptibility to acidic pH as that of *phoPR*-KO. However, all four mycobacterial strains showed comparable growth at pH 7.0. Next, to compare growth of WT-Rv0805 and WT under oxidative stress, cells were grown in presence of increasing concentrations of diamide, a thiol-specific oxidant, and examined by microplate-based Alamar Blue assays (Figure 4 - figure supplement 1A). We have recently shown that *phoPR*-KO is significantly more sensitive to diamide relative to WT (Goar *et al*., 2022). Here, we uncovered that similar to *phoPR*-KO, WT-Rv0805, but not WT-Rv0805M was significantly more susceptible to diamide stress as compared to WT (Fig. 4B). A previous study had reported that *phoP*-deleted mutant strain was more sensitive to Cumene Hydrogen Peroxide (CHP), suggesting a role of PhoP in regulating mycobacterial stress response to oxidative stress (Walters *et al*., 2006). To compare sensitivity to CHP, we grew mycobacterial strains in presence of 50 µM CHP for 24 hours and determined their survival by enumerating CFU values (Fig. 4C). In this case we were unable to perform Alamar Blue-based survival assays requiring a longer time because of the bactericidal property of CHP. Our CFU data highlight that WT-Rv0805, but not WT-Rv0805M, displayed a significantly higher growth inhibition relative to WT in the presence of CHP. Together, these results reveal similar behaviour of *phoPR*-KO, and WT-Rv0805 by demonstrating a comparably higher susceptibility of these strains to acidic pH and oxidative stress relative to WT bacteria and indicate a link between intra-mycobacterial cAMP level and bacterial stress response. It appears that at least one of the mechanisms by which PhoP contributes to global stress response is attributable to maintenance of cAMP level.

**Figure 4:**
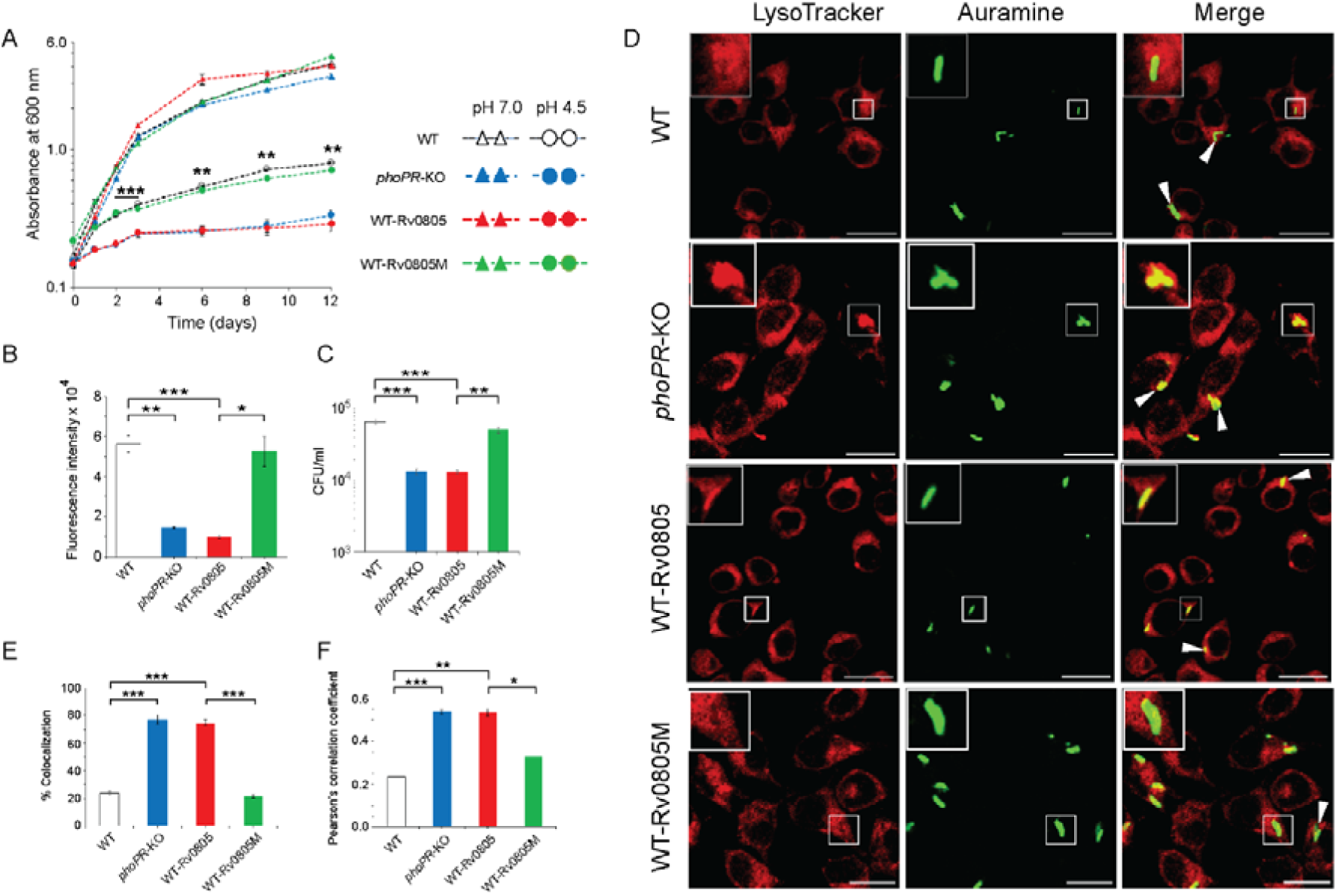
Regulation of cAMP level and its effect on mycobacterial stress tolerance and survival in macrophages. (A) To compare susceptibility to low pH conditions, indicated mycobacterial strains were grown at pH 4.5, and similar to *phoPR*-KO (grey circles), WT-Rv0805 (red circles) shows a significant growth defect relative to WT (empty circles). However, WT-Rv0805M (green circles) grows comparably well as that of the WT (empty circles). In contrast, at pH 7.0 all four mycobacterial strains (WT, empty triangles; *phoPR*-KO, grey triangles; WT-Rv0805, red triangles; WT-Rv0805M, green triangles) displayed comparable growth. (B) Microplate-based assay using Alamar Blue was utilized to examine mycobacterial sensitivity to increasing concentrations of diamide. In this assay, reduction of Alamar Blue correlates with the change of a non-fluorescent blue to a fluorescent pink appearance, which is directly proportional to bacterial growth. Survival of indicated mycobacterial strains, under normal conditions and in presence of 5 mM diamide, were determined by plotting fluorescence intensity. The data is normalized relative to WT grown in presence of 5 mM diamide. (C) To compare susceptibility to stress conditions, these mycobacterial strains were next grown in the presence of 50 µM Cumene Hydrogen Peroxide (CHP). In presence of CHP, WT-Rv0805 (red column) but not WT-Rv0805M (green column), shows a significant growth defect [relative to WT (empty column)] in striking similarity to *phoPR*-KO (grey column). Note that similar to *phoPR*-KO, WT-Rv0805 shows a comparably higher sensitivity to CHP relative to WT bacilli. However, WT-Rv0805M expressing a mutant Rv0805, shows a significantly lower sensitivity to CHP relative to WT-Rv0805, as measured by the corresponding CFU values. The growth experiments were performed in biological duplicates, each with two technical replicates (**P≤0.01; ***P≤0.001). (D) Murine macrophages were infected with indicated *M. tuberculosis* H37Rv strains. The cellular organelle was made visible by LysoTracker; mycobacterial strains were stained with phenolic auramine solution, and the confocal images display merge of two fluorescence signals (Lyso Tracker: red; H37Rv: green; scale bar: 10 µm). The insets in the merge panels indicate bacteria which either have inhibited or facilitated trafficking into lysosomes. <u>White *arrowheads* in the merge panels indicate non-colocalization which remains higher in WT-H37Rv and WT-Rv0805M relative to *phoPR*-KO or WT-Rv0805.</u> (E) Bacterial co-localization of *M. tuberculosis* H37Rv strains. The percentage of auramine labelled strains co-localized with Lysotracker was determined by counting at least 100 infected cells in 10 different fields. The results show the average values with standard deviation determined from three independent experiments (***P≤ 0.001). (F) Pearson’s correlation coefficient of images of internalized auramine-labelled mycobacteria and Lysotracker red marker in RAW 264.7 macrophages. Data are representative of mean ± S.D., derived from three independent experiments (*P<0.05; ***P<0.001).

A previous study showed that *rv0805* over-expression in *M. smegmatis* influences cell wall permeability (Podobnik *et al*, 2009). Having shown a significantly higher sensitivity of WT-Rv0805 to low pH and oxidative stress (relative to WT), we sought to investigate whether altered cell wall structure/properties of the mycobacterial strain contribute to elevated stress sensitivity. We compared expression level of lipid biosynthetic genes, which encode part of cell-wall structure of the bacilli (Gonzalo Asensio *et al*., 2006a; Walters *et al*., 2006). Our results suggest that in contrast to *phoPR*-KO, both WT-Rv0805 and WT-Rv0805M share a comparable expression profile of complex lipid biosynthesis genes as that of WT (Figure 4 - figure supplement 1B). These results suggest that both strains expressing wild type or mutant PDEs share a largely similar cell-wall properties and are consistent with (a) a recent study reporting no significant effect of cAMP dysregulation on mycobacterial cell wall structure/permeability (Wong *et al*, 2023), and (b) role of PhoP in cell wall composition and complex lipid biosynthesis (Gonzalo Asensio *et al*., 2006a; Goyal *et al*., 2011; Walters *et al*., 2006). These results support our view that higher susceptibility of WT-Rv0805 to stress conditions, is attributable to its reduced cAMP level.

To investigate the impact of mycobacterial cAMP level *in vivo*, we studied infection of murine macrophages using WT, WT-Rv0805, and WT-Rv0805M (Fig. 4D). In this assay, WT-H37Rv inhibits phagosome maturation, whereas phagosomes with *phoPR*-KO mature into phagolysosomes (Anil Kumar *et al*., 2016). In our present experimental set up, although WT bacilli inhibited phagosome maturation, infection of macrophages with WT-Rv0805 and *phoPR*-KO matured into phagolysosomes, suggesting increased trafficking of the bacilli to lysosomes. Under identical conditions, WT-Rv0805M could effectively inhibit phagosome maturation just as WT bacteria. Results from co-localization experiments are plotted as Fig. 4E and as Pearson’s correlation coefficient of the quantified co-localization signals as Fig. 4F. These data suggest reduced ability of WT-Rv0805, but not WT-Rv0805M (relative to WT) to inhibit phagosome maturation. From these results, we suggest that ectopic expression of *rv0805* impacts phagosome maturation arguing in favour of a role of PhoP in influencing phagosome-lysosome fusion in macrophages.

### Intra-bacterial cAMP level and its effect on *in vivo* survival of mycobacteria

To examine the effect of intra-bacterial cAMP level on *in vivo* survival, mice were infected with mycobacterial strains via the aerosol route. Day 1 post-infection, CFU analyses revealed a comparable count of four mycobacterial strains (∼100 bacilli) in the mice lungs. However, for WT-Rv0805, the CFU recovered from infected lungs 4 weeks post-infection declined by ∼218-fold relative to the lungs infected with WT bacteria (Fig. 5A). In contrast, the CFU recovered from infected lungs after 4 weeks of infection by WT-Rv0805M marginally declined by ∼7-fold relative to the lungs infected with WT bacilli. These results suggest a significantly compromised ability of WT-Rv0805 (relative to WT) to replicate in the mice lungs. Note that *phoPR*-KO, under the conditions examined, showed a ∼ 246-fold lower lung burden compared to WT. In keeping with these results, while the WT bacilli disseminated to the spleens of infected mice, a significantly lower count of WT-Rv0805 was recovered from the spleens after 4 weeks of infection (Fig. 5B). Thus, we suggest that one of the reasons which accounts for an attenuated phenotype of *phoPR*-KO in both cellular and animal models is attributable to PhoP-dependent repression of *rv0805* PDE activity, which controls mycobacterial cAMP level.

**Figure 5:**
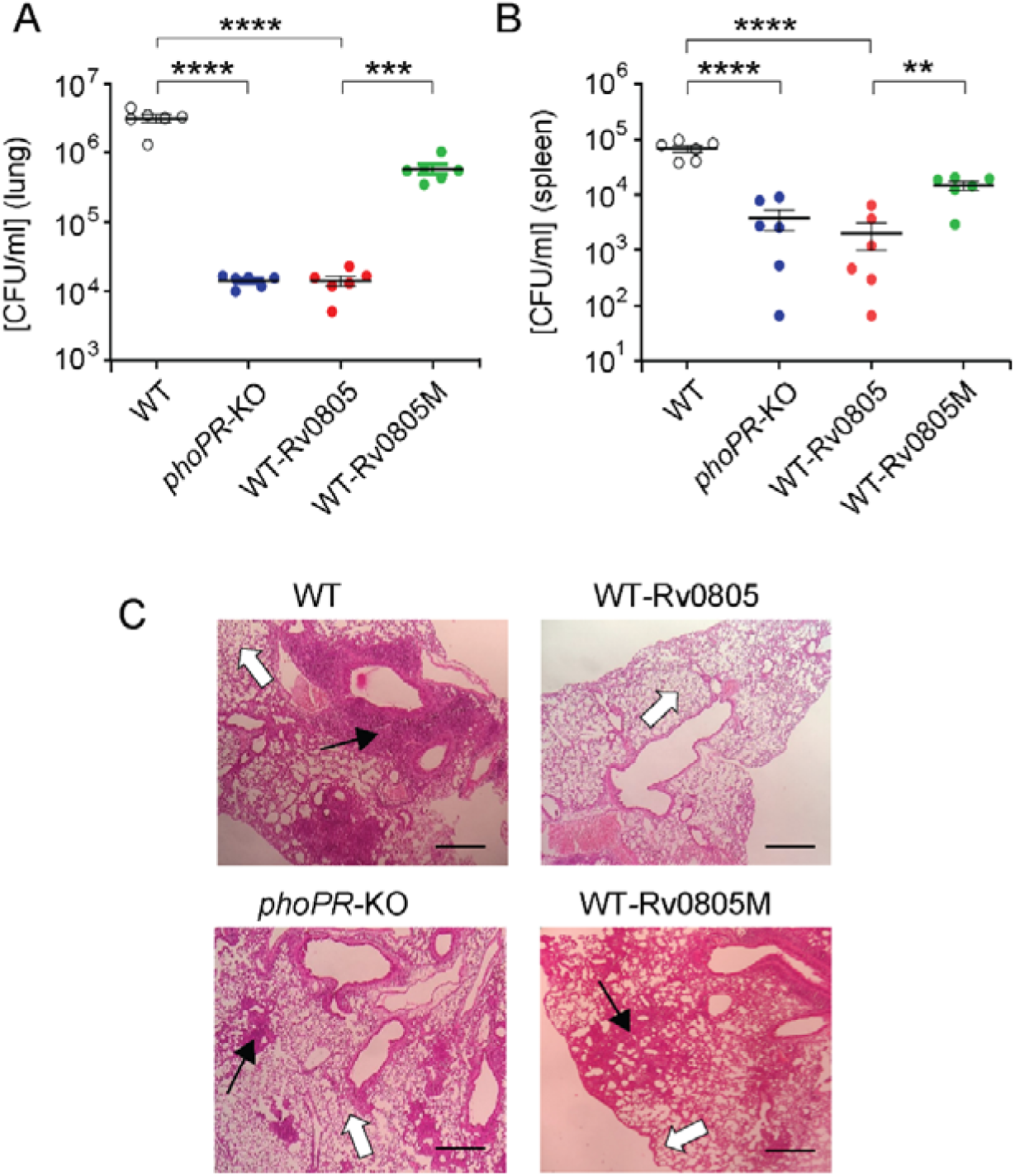
Dysregulation of mycobacterial cAMP level impacts mycobacterial survival *in vivo*. (A-B) Survival of mycobacterial strains in mice (A) lung, and (B) spleen after animals were given an aerosol infection with ∼100 CFU / lung. Mycobacterial load represents mean CFU values with standard deviations obtained from at least five animals per strains used (**P< 0.01; ***P<0.001; ****P<0.0001). (C) Histopathology of lung sections after 4 weeks of infection with indicated bacterial strains. Sections were stained with hematoxylin and eosin, observed under a light microscope, and images of the pathology sections collected at x40 magnification display granulomas (*filled arrows*) and alveolar space (*empty arrows*) (scale bar, 200 µm).

*M. tuberculosis* H37Rv persists within granulomas where it is protected from the anti-mycobacterial immune effectors of the host. Histopathological evaluations showed that WT bacilli - infected lung sections displayed aggregation of granulocytes within alveolar spaces which degenerate progressively to necrotic cellular debris. In contrast, *phoPR*-KO and WT-Rv0805 showed less severe pathology as indicated by decreased tissue consolidation, smaller granulomas, and open alveolar space (Fig. 5C). Together, these results suggest that ectopic expression of *rv0805* in WT bacilli is phenotypically equivalent to deletion of *phoP*, suggesting that failure to maintain cAMP level most likely accounts for attenuated phenotype of the bacilli and absence of immunopathology in the lungs of infected mice.

## Discussion

A number of studies suggest that conditions associated with host environment like low pH, and macrophage interactions often influence mycobacterial cAMP levels (Bai *et al*., 2009; Gazdik & McDonough, 2005). Although many bacterial pathogens modulate host cell cAMP levels as a common strategy, the mechanism of regulation of mycobacterial cAMP-level remains unknown. In this study, we sought to investigate whether PhoP, a master regulator implicated in controlling diverse mycobacterial stress response, regulates mycobacterial cAMP level. We find that under normal conditions as well as under carefully controlled single stress conditions, *phoPR*-KO shows a significantly lower level of cAMP relative to the WT bacilli (Fig. 1A), and complementation of the mutant restored cAMP level. To investigate the mechanism, we next probed regulation of ACs and PDEs (Fig. 2) and demonstrated that PhoP functions as a major repressor of *rv0805*, encoding cAMP specific PDE. Indeed, this newly-identified *rv0805* regulation, coupled with a recent discovery that phosphodiesterase activity of Rv0805 controls propionate detoxification (McDowell *et al*, 2023), fits well with and explains, the previously puzzling *in vivo* observation by Abramovitch et al (Abramovitch *et al*., 2011) that PhoP -controlled *aprABC* locus is associated with the regulation of genes of carbon and propionate metabolism.

Although a large number of ACs are present in *M. tuberculosis* genome, a class III metallo-phosphoesterase Rv0805 was earlier considered the only PDE, specific for mycobacterial cAMP. However, a recent study has identified an atypical class II PDE Rv1339, which upon overexpression reduces cAMP level and contributes to antibiotic sensitivity (Thomson *et al*., 2022). While, the functional role of Rv1339 in *M. tuberculosis* is yet to be understood, crystal structure and biochemical evidence suggest that dimeric Rv0805 is stabilized by the presence of a divalent cation, and remains catalytically active on a broad range of linear and cyclic PDE substrates *in vitro* (Keppetipola & Shuman, 2008; Shenoy *et al*., 2007; Shenoy *et al*., 2005). More recently, cyclic nucleotide hydrolytic activity of mycobacterial Rv0805 has been implicated in propionate detoxification (McDowell *et al*., 2023). However, the mechanism of regulation of Rv0805 and its effect on mycobacterial cAMP level remained unknown before the present study.

To examine biological significance of PhoP-dependent Rv0805 expression, we studied *rv0805* expression under acidic conditions of growth as *phoPR* system is induced under acidic pH both *in vitro* and in macrophages (Abramovitch *et al*., 2011; Bansal *et al*., 2017). A significantly higher repression of *rv0805* expression under acidic pH relative to normal conditions is consistent with activation of PhoP and subsequent repression of *rv0805* (Fig. 3). These results further suggest that effective mitigation of stress by mycobacteria possibly requires a higher cAMP level for survival under intra-phagosomal environment. In keeping with these results, we find that (a) P∼PhoP binds to *rv0805* regulatory region (Fig. 2-figure supplement 1), and (b) PhoP-dependent *rv0805* expression requires PhoR (Figs. 3A-B), the cognate kinase which activates PhoP in a signal-dependent manner (Gupta *et al*., 2006; Singh *et al*, 2023). These results account for a consistently lower level of cAMP in *phoPR*-KO relative to the WT bacilli. Notably, except recently reported PDE Rv1339, Rv0805 has been known as the only cAMP-specific PDE present in the slow growing pathogenic mycobacteria and its closely related species (Matange, 2015; Shenoy *et al*., 2007; Shenoy *et al*., 2005), and Rv1339 expression does not appear to be regulated by the *phoP* locus (Fig. 2). Thus, the above results showing PhoP-dependent repression of *rv0805* activity likely <u>represent the</u> most critical step of regulation of mycobacterial cAMP level <u>under stress</u>. In this connection, our recent results that PhoP interacts with cAMP receptor protein, CRP and a complex of two interacting regulators control expression of virulence determinants (Khan *et al*, 2022), invite speculation of a complex regulatory control of cAMP-responsive mycobacterial physiology.

As one might argue that PhoP deletion and *rv0805* over-expression could be unrelated and independent events, we constructed *phoP* and *rv0805* knock-down mutants to further investigate the PhoP-Rv0805-cAMP pathway. Our objective was to probe regulation of expression (Fig. 3D) and examine the impact on mycobacterial cAMP (Fig. 3E). *phoP*-KD significantly elevated *rv0805* expression; however, *phoP* expression remains unaffected in *rv0805*-KD (Fig. 3D). While elevated *rv0805* expression in *phoP*-KD reduces cAMP level, understandably cAMP level is elevated in *rv0805*-KD mutant (Fig. 3E). These results integrate PhoP-dependent *rv0805* repression with mycobacterial cAMP level, suggesting how *phoP*-deletion or knock-down, at least in part, mimics Rv0805 over-expression. These considerations take on more significance given the fact that these two events have similar consequences on relevant strains with respect to stress tolerance, and survival in cellular and animal models. Thus, our results suggest that ectopic expression of *rv0805* is functionally equivalent to deletion of the *phoP* locus. This observation is in apparent conflict with a previous work by Matange and collaborators (Matange *et al*., 2013), suggesting a cAMP-independent transcriptional response in *rv0805* over-expressing *M. tuberculosis* H37Rv. Although both studies were performed with *rv0805* over-expressing bacilli, the fact that important differences in the expression of PDEs in this study (Matange *et al*., 2013) and in our assays – yielding significantly different levels of *rv0805* expression - most likely account for this discrepancy. While we cannot completely rule out the possibility of cleavage of other cyclic nucleotide(s) by Rv0805 (Keppetipola & Shuman, 2008; Shenoy *et al*., 2007; Shenoy *et al*., 2005) impacting our results, consistent with a previous study our results correlate *rv0805* expression with intra-mycobacterial cAMP level (Agarwal *et al*., 2009).

Further, our data on the effect of expression of cyclic nucleotide-specific PDE Rv0805 or its inactive mutant (Rv0805M) correlate well with enzyme activities of the corresponding PDEs on mycobacterial cAMP levels (Fig. 3C). Thus, we infer that PhoP-dependent regulation of *rv0805* is a critical regulator of intra-mycobacterial cAMP level.

Our experiments to understand physiological significance of PhoP-dependent repression of *rv0805* expression uncovers a comparable stress tolerance of WT-Rv0805 and *phoPR*-KO (significantly reduced relative to WT) (Fig. 4). These results are consistent with the notion that cAMP level, at least in part, accounts for mycobacterial stress response. Along the line, WT-Rv0805 displayed a reduced ability to inhibit phagosome-lysosome fusion like *phoPR-*KO (Fig. 4). Further, we show that WT-Rv0805, unlike the WT bacilli or WT-Rv0805M, shows a significantly reduced intracellular growth in mice as that of *phoPR*-KO (Fig. 5). Thus, these results are of fundamental significance to establish that PhoP contributes to maintenance of cAMP level and integrates it to mechanisms of mycobacterial stress tolerance and intracellular survival. Together, we identify a novel mycobacterial pathway as a therapeutic target and provide yet another example of an intimate link between bacterial physiology and intracellular survival of the tubercle bacilli.

The results reported here are presented schematically in Fig. 6. In summary, upon sensing low acidic pH as a signal PhoR activates PhoP, P∼PhoP binds to *rv0805* upstream regulatory region and functions as a specific repressor of Rv0805. Therefore, we observed (a) a reproducibly lower level of cAMP in *phoPR*-KO relative to WT-H37Rv, (b) a significantly reduced expression of *rv0805* in WT-H37Rv, grown under acidic pH relative to normal conditions, and (c) comparable cAMP levels in *phoPR*-KO and WT-Rv0805. This is why the two strains remain ineffective to mount an appropriate stress response, most likely due to their inability to coordinate regulation of gene expression because of dysregulation of intra-mycobacterial cAMP level. However, without uncoupling regulatory control of PhoPR and *rv0805* expression, we cannot confirm that dysregulation of cAMP level accounts for virulence attenuation of *phoPR*-KO. Given the fact that *rv0805*-depleted *M. tuberculosis* is growth attenuated *in vivo* (McDowell et al., 2023), paradoxically ectopic expression of *rv0805* leads to dysregulated metabolic adaptation, thereby resulting in reduced stress tolerance and intracellular survival.

**Figure 6:**
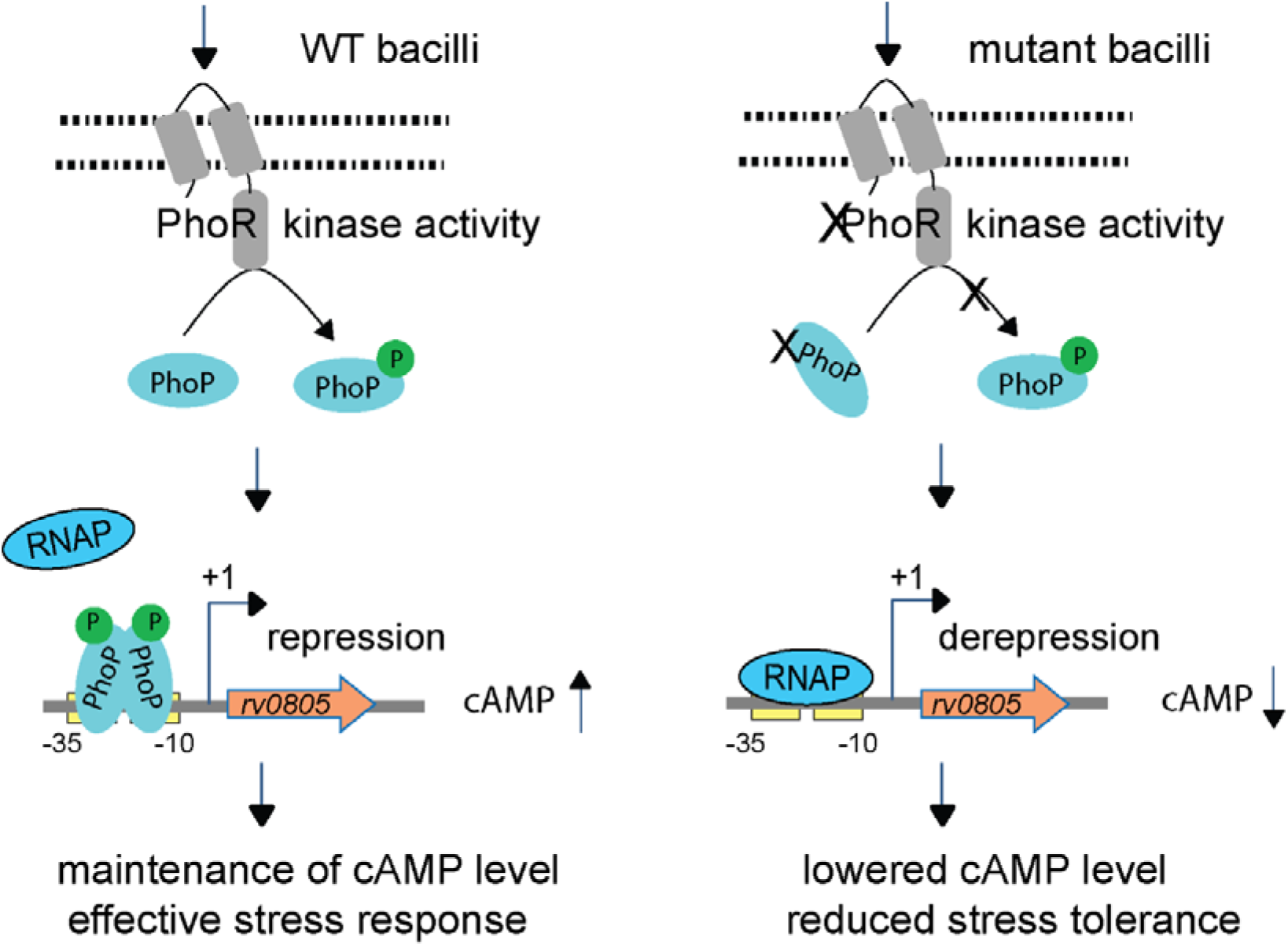
Increased cAMP level and effective stress response versus decreased cAMP level and reduced stress tolerance of mycobacteria. In this model, upon activation by an appropriate signal via the cognate sensor PhoR, P∼PhoP binds to *rv0805* regulatory region and functions as a specific repressor by preventing access for mycobacterial RNA polymerase (RNAP) to bind to the promoter and initiate transcription. In keeping with PhoP-dependent *rv0805* repression, our results demonstrate a reproducibly lower level of cAMP in *phoPR*-KO relative to WT bacilli. Thus, *phoPR*-KO (or WT-Rv0805) remains ineffective to mount an appropriate stress response most likely due to its inability to coordinate regulated gene expression because of dysregulation of cAMP level, accounts for their reduced stress tolerance. Together, these molecular events suggest that failure to maintain cAMP level accounts for attenuated phenotype of the bacilli.

## Materials and Methods

### Bacterial strains and culture conditions

*E. coli* DH5α was used for cloning experiments. *E. coli* BL21(DE3), an *E. coli* B strain lysogenized with λDE3, a prophage expressing T7 RNA polymerase from the IPTG (isopropyl-β-D-thiogalactopyranoside)-inducible *lac*UV5 promoter (Studier & Moffatt, 1986) was used as the host for over-expression of recombinant proteins. *E. coli* strains were grown in LB medium at 37°C with shaking, transformed according to standard procedures and the transformants were selected on media containing appropriate antibiotics plates. WT- and mutant *M. tuberculosis* and *M. smegmatis* mc^2^155 were grown at 37°C in Middlebrook 7H9 broth (Difco) containing 0.2% glycerol, 0.05% Tween-80 and 10% ADC (albumin-dextrose-catalase) or on 7H10-agar medium (Difco) containing 0.5% glycerol and 10% OADC (oleic acid-albumin-dextrose-catalase) enrichment. *phoPR* disruption mutant of *M. tuberculosis* H37Rv (*phoPR*-KO, a kind gift of Dr. Issar Smith) was constructed as described (Walters *et al*., 2006). To this end, a kanamycin-resistant cassette from pUC-K4 was inserted into a unique EcoRV site within the coding region of *phoP* gene, and disruption was confirmed by Southern blot analysis of chromosomal DNA isolated from the mutant. Next, purified plasmid DNAs were electroporated into wild-type *M. tuberculosis* strain by standard protocol (Jacobs *et al*, 1991). To complement *phoPR* expression, pSM607 containing a 3.6-kb DNA fragment of *M. tuberculosis phoPR* including 200-bp *phoP* promoter region, a hygromycin resistance cassette, *attP* site and the gene encoding phage L5 integrase, as detailed earlier (Walters *et al*., 2006) was used to transform *phoPR* mutant to integrate at the L5 *attB* site. Growth, transformation of wild-type (WT), *phoPR*-KO, the complemented mutant (Compl.) *M. tuberculosis* and selection of transformants on appropriate antibiotics plates were performed as described (Goyal *et al*., 2011). When appropriate, antibiotics were used at the following concentrations: hygromycin (hyg), 250 µg/ml for *E. coli* or 50 µg/ml for mycobacterial strains; streptomycin (str), 100 µg/ml for *E. coli* or 20 µg/ml for mycobacterial strains; kanamycin (kan), 20 µg/ml for mycobacterial strains. For *in vitro* growth under specific stress conditions, indicated mycobacterial strains were grown to mid log phase (OD_600_ 0.4-0.6) and exposed to different stress conditions. For acid stress, cells were initially grown in 7H9 media, pH7.0 and on attaining mid log phase it was transferred to acidic media (7H9 media, pH 4.5), and grown for further two hours at 37°C. For oxidative stress, cells were grown in presence of 50 µM CHP (Sigma) for 24 hours or indicated diamide concentration(s) for 7 days. For NO stress, cells grown to mid log phase were exposed to 0.5 mM DataNonoate for 40 minutes (Voskuil *et al*, 2003).

### cAMP measurement

Mycobacterial cell pellets were collected and washed with 1x PBS buffer, cells were resuspended in IP buffer (50 mM Tris pH 7.5, 150 mM NaCl, 1 mM EDTA pH 8.0, 1 mM PMSF, 5% glycerol and 1% TritonX 100) and cell lysates (CL) were prepared by lysing the cells in presence of Lysing Matrix B (100 µm silica beads; MP Bio) using a FastPrep-24 bead beater (MP Bio) at a speed setting of 6.0 for 30 seconds. The procedure was repeated for 10 cycles with incubation on ice in between pulses. The supernatant was collected by centrifugation at 13,000 rpm for 10 minutes and filtered through 0.22 µm filter (Millipore). cAMP levels in the cells were determined in a plate reader by using fluorescence-based cAMP detection kit (Abcam) according to the manufacturer’s recommendations and normalized to the total protein present in the samples as determined by a BCA protein estimation kit (Pierce). For secretion studies, each mycobacterial strain was grown in Sauton’s media as described (Anil Kumar *et al*., 2016), comparable counts of bacterial cells were pelleted, resuspended in 2 ml of Sauton’s media in a 6-well plate format for 2 hours at 37°C, and the supernatants (culture filtrates, CF) were collected for cAMP measurements, as described previously (Anil Kumar *et al*., 2016).

### Cloning

*M. tuberculosis* full-length ORFs of interest were cloned between EcoRI and HindIII sites of the mycobacterial expression vector pSTKi (Parikh *et al*., 2013) and expressed from the P_myc1_*tetO* promoter. Mutation in Rv0805 was introduced by two-stage overlap extension method using mutagenic primers (Supplementary file 1b), and the construct was verified by DNA sequencing. For over-expression of WT-or mutant PDEs, WT bacilli was transformed with pST-rv0805 or pST-rv0805M to generate WT-Rv0805 or WT-Rv0805M, respectively.

### EMSA

rv0805up DNA probe was PCR amplified, resolved on an agarose gel, recovered by gel extraction, and end-labelled with [γ-^32^P ATP] (1000 Ci nmol^-1^) using T4 polynucleotide kinase. The end-labelled DNA probe was purified from free label by Sephadex G-50 spin columns (GE Healthcare), and incubated with increasing amounts of purified PhoP in a total volume of 10 µl binding mix (50 mM Tris-HCl, pH 7.5, 50 mM NaCl, 0.2 mg/ml of bovine serum albumin, 10% glycerol, 1 mM dithiothreitol, ≍ 50 ng of labelled DNA probe, and 0.2 µg of sheared herring sperm DNA) at 20°C for 20 minutes. DNA-protein complexes were resolved by electrophoresis on a 6% (w/v) polyacrylamide gel (non-denaturing) in 0.5X TBE (89 mM Tris-base, 89 mM boric acid and 2 mM EDTA) at 70 V and 4°C, and the position of the radioactive material was determined by exposure to a phosphor storage screen.

### Construction of *M. tuberculosis phoP* and *rv0805* knock-down mutants

In this study, we utilized a previously reported CRISPRi system (Singh *et al*., 2016) to construct knock-down mutants of *phoP* and r*v0805* (*phoP*-KD, and *rv0805*-KD, respectively). This approach efficiently inhibits expression of target genes via inducible expression of dCas9 along with gene specific guide RNAs (sgRNA). The RNAs were 20 nt long and complementary to the non-template strand of the target gene. The sgRNAs of *phoP* and *rv0805* were cloned in a vector pRH2521 using BbsI enzyme and the constructs were confirmed by sequencing. The corresponding clones were used to transform *M. tuberculosis* harbouring pRH2502, a vector expressing an inactive version of *Streptococcus pyogenes* cas9 (dcas9). To express dcas9 and repress sgRNA-targeted genes (*phoP* or *rv0805*), the bacterial strains were grown with or without 600 ng/ml of anhydro-tetracycline (ATc) every 48 hours, and cultures were grown for 4 days. RNA isolation was carried out, and RT-qPCR experiments verified significant repression of target genes. For the induced strains (in presence of ATc) expressing sgRNAs targeting +155 to +175 (relative to *phoP* translational start site) and +224 to +244 sequences (relative to *rv0805* translational start site), we obtained approximately 85% and 90% reduction of *phoP* and *rv0805* RNA abundance, respectively, relative to corresponding un-induced strains. The oligonucleotides used to generate gene-specific sgRNA constructs and the plasmids utilized in knock-down experiments are listed in Supplementary file 1b.

### RNA isolation

Total RNA was extracted from exponentially growing bacterial cultures grown with or without specific stress as described above. Briefly, 25 ml of bacterial culture was grown to mid-log phase (OD_600_= 0.4 to 0.6) and combined with 40 ml of 5 M guanidinium thiocyanate solution containing 1% β-mercaptoethanol and 0.5% Tween 80. Cells were pelleted by centrifugation, and lysed by re-suspending in 1 ml Trizol (Ambion) in the presence of Lysing Matrix B (100 µm silica beads; MP Bio) using a FastPrep-24 bead beater (MP Bio) at a speed setting of 6.0 for 30 seconds.

The procedure was repeated for 2-3 cycles with incubation on ice in between pulses. Next, cell lysates were centrifuged at 13000 rpm for 10 minutes; supernatant was collected and processed for RNA isolation using Direct-Zol^TM^ RNA isolation kit (ZYMO). Following extraction, RNA was treated with DNAse I (Promega) to degrade contaminating DNA, and integrity was assessed using a Nanodrop (ND-1000, Spectrophotometer). RNA samples were further checked for intactness of 23S and 16S rRNA using formaldehyde-agarose gel electrophoresis, and Qubit fluorometer (Invitrogen).

### Quantitative Real-Time PCR

cDNA synthesis and PCR reactions were carried out using total RNA extracted from each bacterial culture, and Superscript III platinum-SYBR green one-step qRT-PCR kit (Invitrogen) with appropriate primer pairs (2 µM) using an ABI real-time PCR detection system. Oligonucleotide primer sequences used in RT-qPCR experiments are listed in Supplementary file 1a. Control reactions with platinum Taq DNA polymerase (Invitrogen) confirmed absence of genomic DNA in all our RNA preparations, and endogenously expressed *M. tuberculosis rpoB* was used as an internal control. Fold difference in gene expression was calculated using ΔΔC_T_ method (Schmittgen & Livak, 2008). Average fold differences in mRNA levels were determined from at least two biological repeats each with two technical repeats. Non-significant difference is not indicated.

### ChIP assays

ChIP experiments in actively growing cultures of *M. tuberculosis* were carried out essentially as described previously (Fol *et al*, 2006). Immunoprecipitation (IP) was performed using anti-PhoP antibody and protein A/G agarose beads (Pierce). qPCR reactions included PAGE purified primer pairs (Supplementary file 1a) spanning specific promoter regions using suitable dilutions of immunoprecipitated (IP) DNA in a reaction buffer containing SYBR green mix, and one unit of Platinum Taq DNA polymerase (Invitrogen). An IP experiment without adding antibody (mock) was used as the negative control, and data was analysed relative to PCR signal from the mock sample. PCR amplifications were carried out for 40 cycles using serially diluted DNA samples (mock, IP treated and total input) in a real-time PCR detection system (Applied Biosystems). In all cases melting curve analysis confirmed amplification of a single product.

### Immunoblotting

Cell lysates or culture filtrates were resolved by 12% SDS-PAGE and visualized by Western blot analysis using appropriate antibodies. Briefly, resolved samples were electroblotted onto polyvinyl difluoride (PVDF) membranes (Millipore, USA) and were detected by anti-GroEL2 antibody (Sigma), anti CFP-10 antibody (Abcam) or affinity-purified anti-PhoP antibody elicited in rabbit (Alpha Omega Sciences, India). Goat anti-rabbit secondary antibody conjugated to horseradish peroxidase was used, and blots were developed with Luminata Forte Chemiluminescence reagent (Millipore, USA). RNA polymerase was used as a loading control and was detected with monoclonal antibody against β-subunit of RNA polymerase, RpoB (Abcam).

### Alamar Blue assay

In this assay, reduction of Alamar Blue correlates with the change of a non-fluorescent blue to a fluorescent pink appearance, which is directly linked to bacterial growth. *M. tuberculosis* H37Rv was grown in 7H9 media (Difco) with 10% ADS (albumin, dextrose and NaCl) to an OD_600_ of 0.4, and freshly diluted to OD_600_ of 0.02. Next, increasing concentrations of diamide was added to the wells of a 96 -well plate containing 0.05 ml 7H9 media followed by addition of 0.05 ml of *M. tuberculosis* H37Rv culture (0.02 OD_600_). The plate was incubated at 37°C for 7 days.

Finally, 0.02 ml of 0.02% Resazurin (sodium salt, MP Bio), prepared in sterile 7H9 media was added to each of the wells and the change in colour was examined after incubation at 37°C for 16 hours. The fluorescence excitation was at 530 nm and emission was recorded at 590 nm. Efficiency of inhibition was calculated relative to control wells which did not include diamide, and Rifampicin was included as a positive control to confirm validity of the assay.

### Macrophage Infections

Virulence of indicated H37Rv strains were assessed in murine macrophages according to the previously published protocol (Solans *et al*., 2014). Briefly, RAW264.7 macrophages were grown in DMEM media containing 10% Fetal bovine serum at 37°C under 5% CO_2_, and seeded onto #1 thickness, 18 mm diameter glass coverslips in a 12-well plate at a 40% confluency (0.5 million cells). Cells were independently infected with titrated cultures of WT, WT-Rv0805, WT- Rv0805M and *phoPR*-KO strains at a multiplicity of infection (MOI) of 1:5 for 3 hours at 37°C in 5% CO_2_, followed by 1X PBS washes thrice. The macrophages were further incubated for 3 hours at 37°C. After infection, extracellular bacteria were removed by washing thrice with PBS. To visualize trafficking of the tubercle bacilli, mycobacterial strains were stained with phenolic auramine solution (which selectively binds to mycolic acids) for 15 minutes followed by washing with acid alcohol solution and finally with 1X PBS. The cells were stained with 150 nM LysoTracker Red DND-99 (Invitrogen) for 30 minutes in a CO_2_ incubator. Next, the cells were fixed with 4% paraformaldehyde for 15 minutes, washed thrice with PBS, the coverslips were mounted in Slow Fade-Anti-Fade (Invitrogen) and analysed using laser scanning confocal microscope (Nikon) equipped with Argon (488 nm excitation line; 510 nm emission detection) and LD (561 nm excitation line; 594 nm emission detection) laser lines. Digital images were processed with IMARIS imaging software (version 9.20). Details of the experimental methods and the laser/detector settings were optimized using macrophage cells infected with WT-H37Rv as described previously (Anil Kumar *et al*., 2016).

A standard set of intensity threshold was made applicable for all images, and percent bacterial co- localization was determined by analyses of at least 50 infected cells originating from 10 different fields of each of the three independent biological repeats.

### Mouse infections

All experiments pertaining to mice were in accordance with Institutional regulations after review of protocols and approval by the Institutional Animal Ethics Committee (IAEC/17/05, and IAEC/19/02). Mice were maintained and bred in the animal house facility of CSIR-IMTECH. Animal infection studies and subsequent experiments were carried out in the Institutional BSL-3 facility as per institutional biosafety guidelines. Briefly, the experiments were conducted with 8-10 weeks old C57BL/6 mice, infected intranasally and euthanized post-infection for evaluation of bacterial load in lungs and spleens. Infections were given through the respiratory route using an inhalation exposure system (Glass-col) with passaged *M. tuberculosis* H37Rv cultures of mid log phase. The actual bacterial load delivered to the animals was enumerated from 3-5 aerogenically challenged mice, 1day post aerosol challenge. The animals were found to achieve a bacillary deposition of 100 to 200 CFU in the lungs for each strain. Four weeks post infection, the animals were sacrificed by cervical dislocation, lungs and spleens were isolated aseptically from the euthanized animals, homogenized in sterile 1X PBS and plated after serially diluting the lysates on 7H11 agar plates, supplemented with 10% OADC and antibiotics (50 μg/ml carbenicillin, 30 μg/ml polymyxin B, 10 μg/ml vancomycin, 20 μg/ml Trimethoprim, 20 μg/ml cycloheximide, and 20 μg/ml Amphotericin B) to enumerate CFU. For histopathology, left lung lobes were fixed in 10% buffered formalin, embedded in paraffin and stained with haematoxylin and eosin for visualization under the microscope. The level of pathology was scored by analysing perivascular cuffing, leukocyte infiltration, multinucleated giant cell formation and epithelial cell injury.

### Statistical analysis

Data are presented as arithmetic means of the results obtained from multiple replicate experiments ± standard deviations. Statistical significance was determined by Student’s paired t-test using Microsoft Excel or Graph Pad Prism. Statistical significance was considered at P values of 0.05.

## Acknowledgements

We are grateful to G. Marcela Rodriguez and Issar Smith (PHRI, New Jersey Medical School - UMDNJ) for Δ*phoP*-H37Rv, and the complemented *M. tuberculosis* H37Rv strains, Adrie Steyn (University of Alabama) for pUAB300/pUAB400 plasmids, Ashwani Kumar for very helpful discussions, and Sanjeev Khosla for critical reading of the manuscript. We thank members of the Institutional animal facility (iCARE) for their help with approval of our projects from the Institutional Animal Ethics Committee. This study received financial support from intramural grants of CSIR-IMTECH (OLP-0170), CSIR (MLP-0049) and by a research grant (to D.S) from SERB (EMR/2016/004904), Department of Science and Technology (DST). H.K., P.P., H.G., B.B. and N.B. were supported by CSIR pre-doctoral fellowships.

## Supplemental Data

The supplemental data include three (3) supplemental figures, and two supplemental tables (Supplemental files 1a, and 1b, respectively). All data generated or analysed during this study are included in the manuscript and supporting file; source data files have been provided for all relevant figures.

## Legends to supplementary figures

**Figure 1-figure supplement 1: Defined and indicated stress conditions do not influence *in vitro* growth of mycobacterial strains.** To determine the impact of *in vitro* stress on bacterial growth under conditions when intra-mycobacterial cAMP levels were measured, WT bacilli, *phoPR*-KO, and the complemented mutant (Compl.) were exposed to carefully controlled stress conditions as detailed in the Methods, and CFUs enumerated. The results clearly suggest that bacterial viability under indicated conditions of stress do not impact intra-mycobacterial cAMP level.

**Figure 2-figure supplement 1: Probing *in vitro* DNA binding of PhoP to the *rv0805* regulatory region.** (A) Core binding site of PhoP within the promoter region spanning -150 to +1-bp upstream regulatory region of *rv0805* (rv0805up; relative to ORF start site) was identified by the MEME Suit bioinformatic software using previously-identified consensus PhoP binding sequence (**He and Wang, 2014**). Note that the predicted binding site spans from -110 to -127 relative to the ORF start site (p=0.000726). (B) To verify DNA binding *in vitro*, rv0805up was amplified, end-labelled, and EMSA was carried out using radio-labelled rv0805up for binding of increasing concentrations of PhoP (lanes 2-4) and PhoP, pre-incubated in phosphorylation mix with acetyl-phosphate (AcP) (lanes 5-7), respectively. Lane 1 shows the free probe. The binding mixtures in lanes 2-4, and lanes 5-7 contained indicated proteins at 1, 2 and 4 μM, respectively. The position of the radioactive material was determined by exposure to a phosphor storage screen. *Open* and *filled* arrows indicate free probe and slower-moving complexes with band shifts produced in presence of P∼PhoP, respectively.

**Figure 3-figure supplement 1: Ectopic expression of wild-type and mutant *rv0805* (Rv0805M) in WT bacilli.** (A) Real-time RT-qPCR was carried out to compare expression of indicated phosphodiesterases (PDEs) in WT (empty bar), WT-Rv0805 (red bar) and WT-Rv0805M (green bar), respectively, as described in the Methods. Average fold difference in mRNA levels from two biological repeats (each with a technical repeat) were determined as described in the Methods (**P≤0.01; ***P≤0.001). Nonsignificant difference is not indicated. (B) To compare growth of WT and the variant mycobacterial strains, CFU values were determined under normal conditions of growth as described in the methods.

**Figure 4-figure supplement 1: PhoP-dependent *rv0805* expression contributes to mycobacterial survival under oxidative stress.** (A) In this experiment, we compared metabolic activity of WT, *phoPR*-KO, WT-Rv0805, and WT-Rv0805M, grown in presence of increasing concentrations of diamide by using Alamar Blue assay. Because reduction of Alamar Blue correlates with the change of a non-fluorescent blue to a fluorescent pink appearance, mycobacterial metabolic activity could be assessed by monitoring fluorescence. Two following controls were included in the Alamar Blue assays: Rifampicin control, reflecting mycobacterial growth inhibition to confirm validity of the assay, and no bacteria control indicated at the left top corner of the plate. (B) Next, effect of over expression of *rv0805* on mycobacterial cell -wall structure was assessed by analysing expression of lipid biosynthetic genes of indicated mycobacterial strains by RT-qPCR. Average fold difference in mRNA levels from two biological repeats (each with a technical repeat) were determined as described in the Methods (***P≤0.001). Nonsignificant difference is not indicated.

## Legends to supplementary Tables

Supplementary file 1a

Sequences of oligonucleotide primers used in RT-qPCR and ChIP-qPCR measurements reported in this study

Supplementary file 1b

Sequences of oligonucleotide primers used for amplification and cloning, and plasmids used in this study

## Source data files for supplementary figures

Figure 1-figure supplement 1

Figure 2-figure supplement 1

Figure 3-figure supplement 1

Figure 4-figure supplement 1

## Author contributions

H.K., P.P., H. G., B.B. and D.S. designed research; H.K., P. P., H.G., and B.B. performed research; N.B. contributed analytical tools; H.K., P. P., H. G., B.B., N.B., and D.S. analysed data; and D.S. wrote the manuscript.

## Supplemental Data

The supplemental data include four supplemental figures and two supplemental tables (supplementary files 1a and 1b, respectively). All other data are part of the paper and its supplemental files.

