## Supplemental Information for "*Mycobacterium tuberculosis* PhoP integrates stress response to intracellular survival by regulating cAMP level"

Dibyendu Sarkar^1,4,^ *

^1^CSIR-Institute of Microbial Technology, Sector 39 A, Chandigarh 160036, India

^2^These authors contributed equally to this work.

^3^Present Address: Department of Medicine, Division of Hematology-Oncology

UT Southwestern Medical Center, Dallas, TX 75235

^4^Academy of Scientific and Innovative Research (AcSIR), Ghaziabad, India

^5^Present address: Department of Biosciences and Bioengineering

Indian Institute of Technology Bombay

Powai, Mumbai-400076, India

Running title: Regulation of mycobacterial cAMP

Key Words: cAMP level; Mycobacterium; PhoP; stress response; virulence regulation

*Address correspondence to: Dibyendu Sarkar, CSIR-Institute of Microbial Technology,

**Supplementary File 1a**

Sequences of oligonucleotide primers used in RT-qPCR and ChIP-qPCR measurements

reported in this study

| **Primers^a^** | **Sequences (5’-3’)** | **Reference** |
| --- | --- | --- |
| FPrv0386RT | GGCCCAGATCCTTACCTTTC | This study |
| RPrv0386RT | TGTGCAGCACTTCCTGAGAC | This study |
| FPrv0891cRT | TCAAACGGTACGAGGGTGAT | This study |
| RPrv0891cRT | CACAACTGCATGACCCATTC | This study |
| FPrv1264RT | CAGCTAGGCGAAGTGGTGTC | This study |
| RPrv1264RT | GGGAAAGTTGTTGTCGGTGT | This study |
| FPrv1339RT | CGCCGTTGGTTATCGACTTC | This study |
| RPrv1339RT | AACAGTCGTCAATCTCCCCA | This study |
| FPrv1625cRT | TGAATTTGCCCCACCGAATC | This study |
| RPrv1625cRT | CAGCGCAATTGAAGGATCCA | This study |
| FPrv1647RT | GCCCAAGATGCTGTGAAGTC | This study |
| RPrv1647RT | AACTCACTTTGCGGGATCAG | This study |
| FPrv2488cRT | TTGCTGTTGGCATCATGTCT | This study |
| RPrv2488cRT | CTTCTGGGCATCATCTAGGC | This study |
| FPrv0805RT | GCCGAACTACGCAAATTCTT | This study |
| RPrv0805RT | ATCCAAAACACTCGGAATCG | This study |
| FPrv1357cRT | TCCTCGTCTACCAGCCAATC | This study |
| RPrv1357cRT | GAGACGTTGACGCTGACAAA | This study |
| FPrv2837cRT | AGCAGGACCTTGATGGACAG | This study |
| RPrv2837cRT | GTTCGACCTCCTTGAACACC | This study |
| FPpks2RT | GTTGTGGAAGGCGTTGTTAC | (Goyal *et al*, 2011) |
| RPpks2RT | GTCGTAGAACTCGTCGCAAT | (Goyal *et al.*, 2011) |
| FPmsl3RT | GTGAAAACAAACTTCGGTCAC | (Goyal *et al.*, 2011) |
| RPmsl3RT | ACAAAGAGTTCAGTGTCAATCTCAG | (Goyal *et al.*, 2011) |
| FPlipFRT | TAGTGGCCATCTCTCCGTTG | (Bansal *et al*, 2017) |
| RPlipFRT | AGCGGCTCATAGAGGTCTTC | (Bansal *et al.*, 2017) |
| FP16SrDNART | CTGAGATACGGCCCAGACTC | (Khan *et al*, 2022) |
| RP16SrDNART | CGTCGATGGTGAAAGAGGTT | (Khan *et al.*, 2022) |

^a^FP, forward primer; RP, reverse primer

**Supplementary File 1b**

Sequences of oligonucleotide primers for amplification and cloning, and plasmids used in this study

| **Primers^a^** | **Sequences (5’-3’)** | **Reference** |
| --- | --- | --- |
| FPrv0805up | CGGCGTTCTGGTATCTCG | This study |
| RPrv0805up | TAAGAGAACGTAATCCGG | This study |
| FPrv0805start | AATAATGATATCGTGCATAGACTT | This study |
| RPrv0805stop | AATAATAAGCTTTCAGTCGACGGGA | This study |
| FPrv0805N97A | TGGGTGATGGGTGCACACGACGACCG | This study |
| RPrv0805N97A | CGGTCGTCGTGTGCACCCATCACCCA | This study |
| 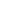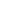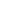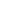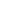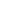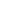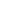FPphoPsg | GGGAGATCCAGCGCCTGTGCCCCG | This Study |
| 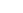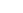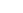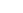RPphoPsg | AAACCGGGGCACAGGCGCTGGATC | This Study |
| FPrv0805sg | GGGAGCTCGACCAGGCCTCGGAGC | This Study |
| 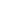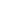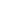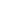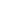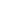RPrv0805sg | AAACGCTCCGAGGCCTGGTCGAGC | This Study |
| FPpRH2521seq | AAACTCTAGAAATATTGGATCG | This study |
| RPpRH2521seq | CCTAATGACCATGGTGACCTC | This study |
| **Plasmids** |  |  |
| p19Kpro^b^ | Mycobacteria expression vector, Hyg^r^ | (De Smet *et al*, 1999) |
| p19Kpro-*phoP* | His_6_-tagged PhoP residues 1-247 cloned in p19Kpro | (Anil Kumar *et al*, 2016) |
| p19Kpro-*phoP*(FLAG) | FLAG-tagged PhoP residues1-247 cloned in p19Kpro | This study |
| pSTKi^c^ | Integrative mycobacterial expression vector, Kan^r^*^c^* | (Parikh *et al*, 2013) |
| pSTKi-*pde* | PDE (*rv0805*) residues 1-957 cloned in pSTKi | This study |
| pSTKi-*pde^M^* | PDE (*rv0805M*) Asn-97 codon mutated to Ala in pSTKi-*pdeM* | This study |
| pRH2502 | Integrative mycobacterial expression vector, Kan^r, c^ | (Singh *et al*, 2016) |
| pRH2521 | Episomal expression vector, Hyg^r,b^ | (Singh *et al.*, 2016) |
| pRH2521-phoPsg | pRH2521 vector expressing *phoP* guide RNA, Hyg^r,b^ | This study |
| pRH2521-rv0805sg | pRH2521 vector expressing *rv0805* guide RNA, Hyg^r,b^ | This study |

^a^FP, forward primer; RP, reverse primer;

^b^ hygromycin resistance (Hyg^r^)

^c^ kanamycin resistance (Kan^r^)

**Figure 1-figure supplement 1**

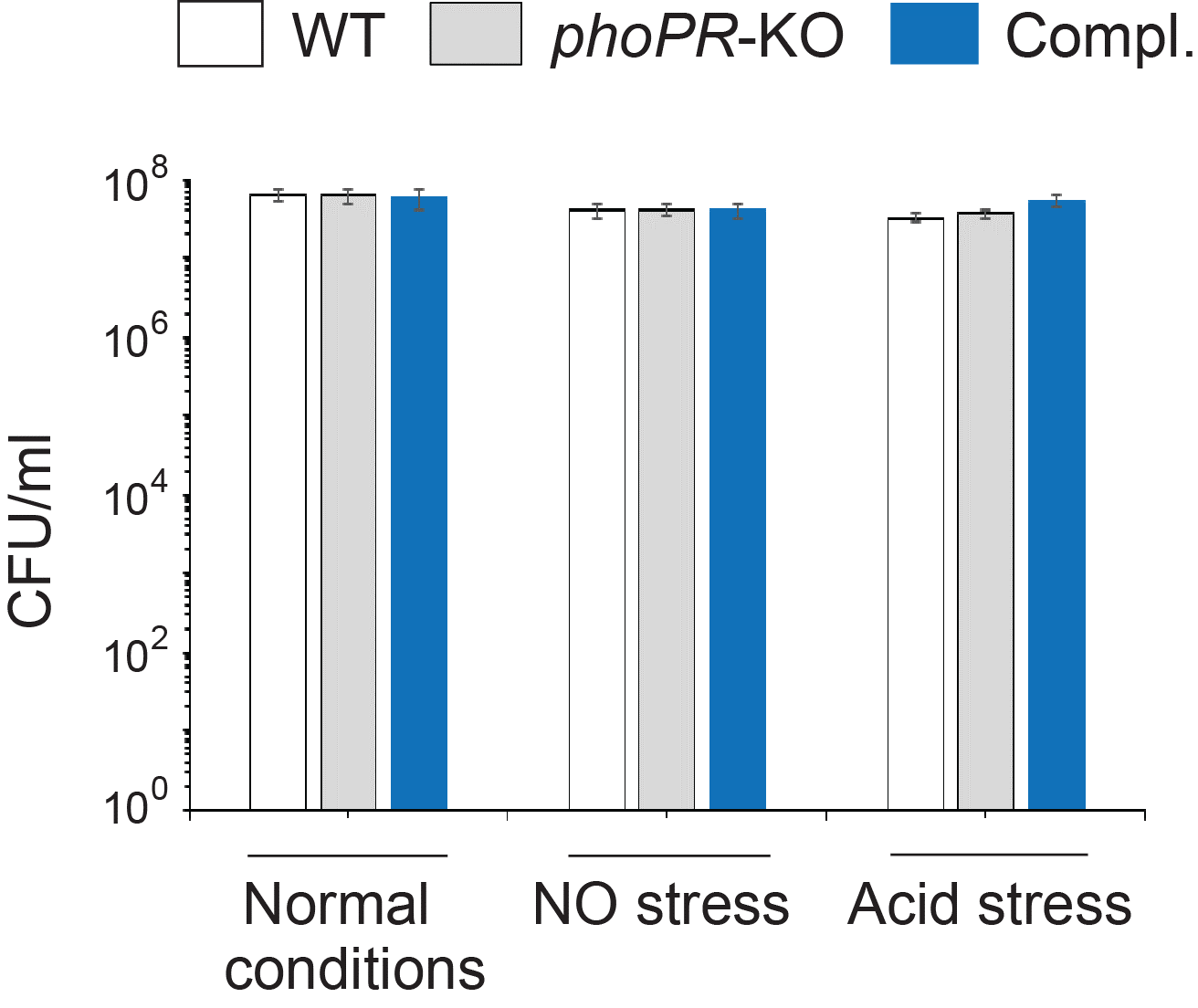

**Figure 1-figure supplement 1: Defined and indicated stress conditions do not influence *in vitro* growth of mycobacterial strains.** To determine the impact of *in vitro* stress on bacterial growth under conditions when intra-mycobacterial cAMP levels were measured, WT bacilli, *phoPR*-KO, and the complemented mutant (Compl.) were exposed to carefully controlled stress conditions as detailed in the Methods, and CFUs enumerated. The results clearly suggest that bacterial viability under indicated conditions of stress do not impact intra-mycobacterial cAMP level.

**Figure 2-figure supplement 1**

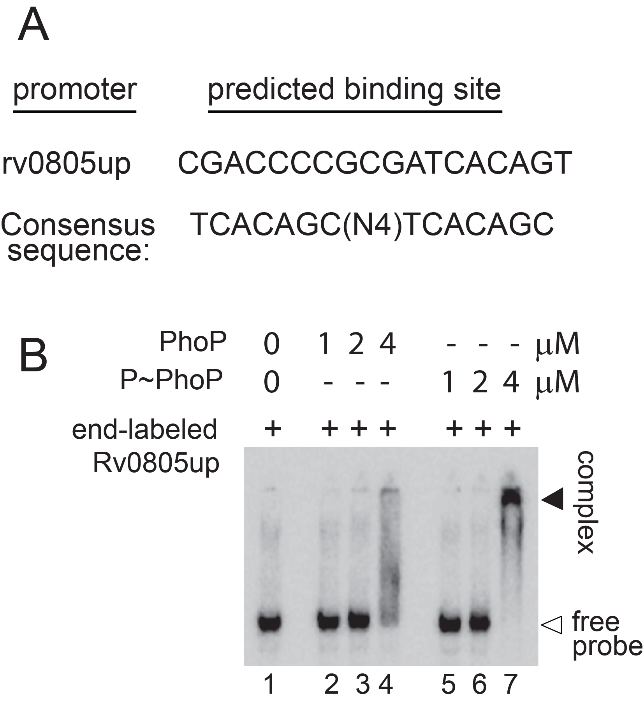

**Figure 2-figure supplement 1:** Probing *in vitro* DNA binding of PhoP to the *rv0805* regulatory region. (A) Core binding site of PhoP within the promoter region spanning -150 to +1-bp upstream regulatory region of *rv0805* (rv0805up; relative to ORF start site) was identified by the MEME Suit bioinformatic software using previously-identified consensus PhoP binding sequence (He & Wang, 2014). Note that the predicted binding site spans from -110 to -127 relative to the ORF start site (p=0.000726). (B) To verify DNA binding *in vitro*, rv0805up was amplified, end-labelled, and EMSA was carried out using radio-labelled rv0805up for binding of increasing concentrations of PhoP (lanes 2-4) and PhoP, pre-incubated in phosphorylation mix with acetyl-phosphate (AcP) (lanes 5-7), respectively. Lane 1 shows the free probe. The binding mixtures in lanes 2-4, and lanes 5-7 contained indicated proteins at 1, 2 and 4 μM, respectively. The position of the radioactive material was determined by exposure to a phosphor storage screen. *Open* and *filled* arrows indicate free probe and slower-moving complexes with band shifts produced in presence of P~PhoP, respectively.

**Figure 3-figure supplement 1**

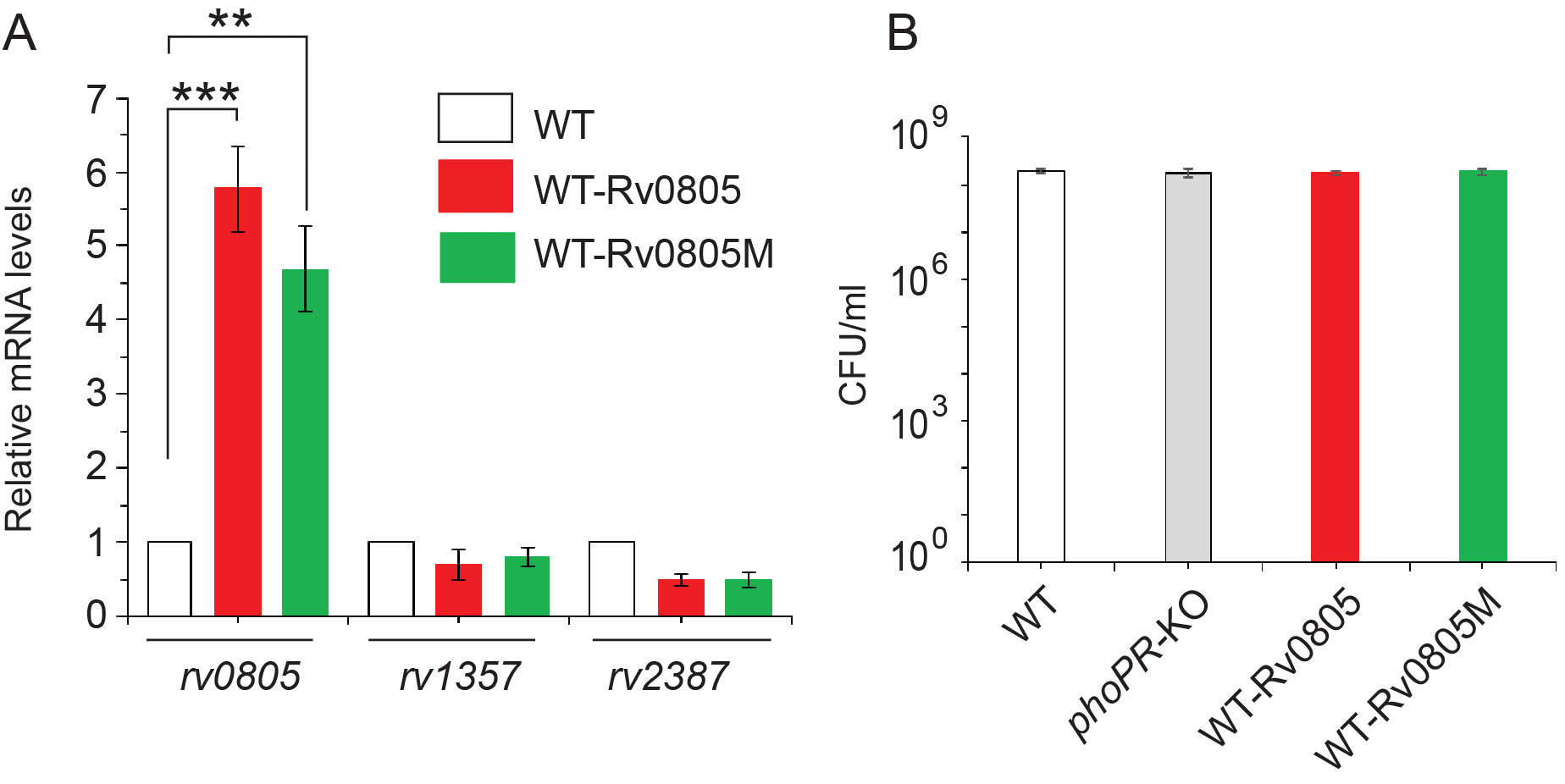

**Figure 3-figure supplement 1: Ectopic expression of WT and mutant *rv0805* (Rv0805M) in WT bacilli.** (A) Real-time RT-qPCR was carried out to compare expression of indicated phosphodiesterases (PDEs) in WT (empty bar), WT-Rv0805 (red bar) and WT- Rv0805M (green bar), respectively, as described in the Methods. Average fold difference in mRNA levels from two biological repeats (each with a technical repeat) were determined as described in the Methods (**P≤0.01; ***P≤0.001). Nonsignificant difference is not indicated. (B) To compare growth of WT and the variant mycobacterial strains, CFU values were determined under normal conditions of growth as described in the methods.

**Figure 4-figure supplement 1**

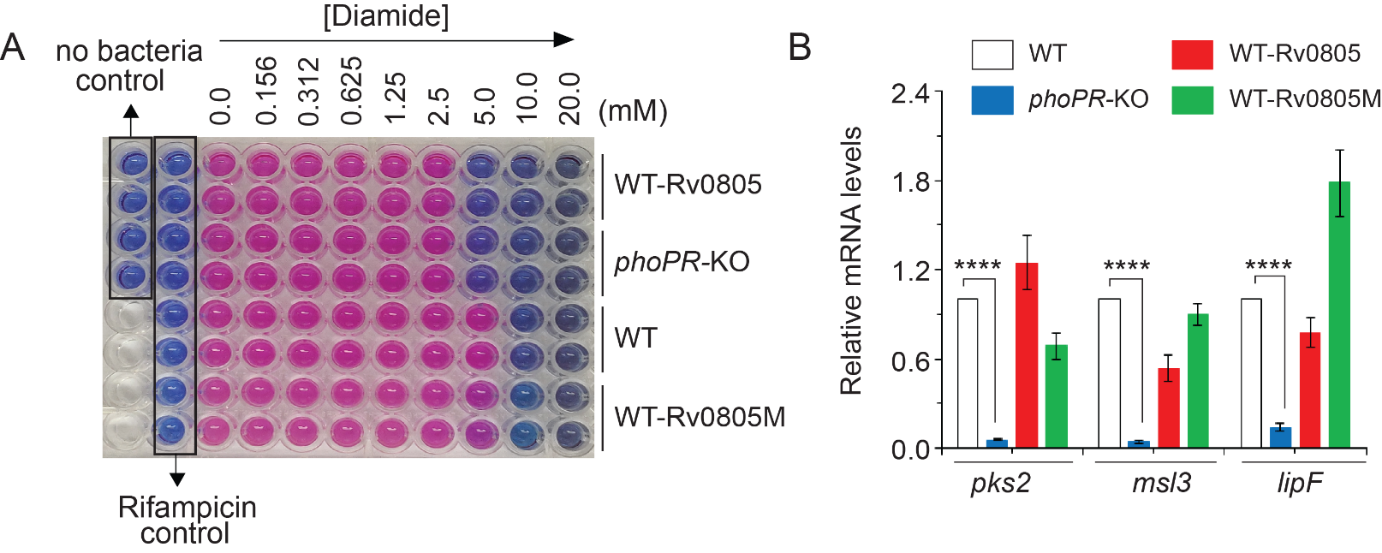

**Figure 4-figure supplement 1: PhoP-dependent *rv0805* expression contributes to mycobacterial survival under oxidative stress.** (A) In this experiment, we compared metabolic activity of WT, *phoPR*-KO, WT-Rv0805, and WT-Rv0805M, grown in presence of increasing concentrations of diamide by using Alamar Blue assay. Because reduction of Alamar Blue correlates with the change of a non-fluorescent blue to a fluorescent pink appearance, mycobacterial metabolic activity could be assessed by monitoring fluorescence. Two following controls were included in the Alamar Blue assays: Rifampicin control, reflecting mycobacterial growth inhibition to confirm validity of the assay, and no bacteria control indicated at the left top corner of the plate. (B) Next, effect of over expression of *rv0805* on mycobacterial cell -wall structure was assessed by analysing expression of lipid biosynthetic genes of indicated mycobacterial strains by RT-qPCR. Average fold difference in mRNA levels from two biological repeats (each with a technical repeat) were determined as described in the Methods (***P≤0.001). Nonsignificant difference is not indicated.
